# ATM and PRDM9 Regulate SPO11-bound Recombination Intermediates During Meiosis

**DOI:** 10.1101/2019.12.18.881409

**Authors:** Jacob Paiano, Wei Wu, Shintaro Yamada, Nicholas Sciascia, Elsa Callen, Ana Paola Cotrim, Rajashree A. Deshpande, Yaakov Maman, Amanda Day, Tanya T. Paull, André Nussenzweig

## Abstract

Meiotic recombination is initiated by genome-wide SPO11-induced double-strand breaks (DSBs) that are processed by MRE11-mediated release of SPO11. The DSB is then resected and loaded with DMC1/RAD51 filaments that invade homologous chromosome templates. In most mammals, DSB locations (“hotspots”) are determined by the DNA sequence specificity of PRDM9. Here, we demonstrate the first direct detection of meiotic DSBs and resection in vertebrates by performing END-seq on mouse spermatocytes using low sample input. We find that DMC1 limits both the minimum and maximum lengths of resected DNA, whereas 53BP1, BRCA1 and EXO1 play surprisingly minimal roles in meiotic resection. Through enzymatic modifications to the END-seq protocol that mimic the *in vivo* processing of SPO11, we identify a novel meiotic recombination intermediate (“SPO11-RI”) present at all hotspots. The SPO11-bound intermediate is dependent on PRDM9 and caps the 3’ resected end during engagement with the homologous template. We propose that SPO11-RI is generated because chromatin-bound PRDM9 asymmetrically blocks MRE11 from releasing SPO11. In *Atm^−/−^* spermatocytes, SPO11-RI is reduced while unresected DNA-bound SPO11 accumulate because of defective MRE11 initiation. Thus in addition to their global roles in governing SPO11 breakage, ATM and PRDM9 are critical local regulators of mammalian SPO11 processing.

## Main

Recombination between homologous chromosomes during meiosis requires DNA double-strand break (DSB) formation by the topoisomerase-like protein SPO11 ^1^. After cutting, SPO11 remains covalently bound to a two-nucleotide, 5’ overhang at both ends of the DNA via phosphotyrosyl linkage. Recombination then begins with the processing of SPO11-bound DSBs into resected 3’ single-stranded DNA (ssDNA) tails that preferentially invade the homologous chromosome by the recombinases DMC1 and RAD51. Studies in budding yeast *Saccharomyces cerevisiae* determined that the MRE11/RAD50/NBS1 (MRN) complex detects SPO11 and cooperates with Sae2 to produce a nick on the SPO11-bound strand via MRE11 endonuclease activity ^2^. The nick serves as an entry point for both short-range MRE11 3’-5’ exonuclease activity to degrade back to the DSB, thereby removing covalently bound SPO11 attached to a ssDNA oligonucleotide, as well as for more extensive long-range processing of 5’ strands (**Extended Data Fig. 1a**) ^2^. In budding yeast, Exo1 nuclease is uniquely responsible for this long-range 5’-3’ resection ^3^. Moreover, short- and long-range resection are tightly coupled in a single processive reaction (**Extended Data Fig. 1a**). As a result, meiotic DSBs are maximally resected as soon as they appear and unresected SPO11-bound DSBs are extremely rare ^4–6^. While ATM has been shown to regulate DSB numbers and locations ^7, 8^, its remains unclear whether it also functions downstream in regulating SPO11 processing and resection.

Distinct from yeast, DSB hotspots in mice and humans are determined by the DNA binding specificity of the PRDM9 methyltransferase ^9^. Besides positioning DSBs, PRDM9 binding activity also reorganizes nucleosomes in a manner that creates a nucleosome depleted region (NDR) within which DSBs and PRDM9 itself are centered ^10^. Moreover, PRDM9 has been suggested to have a role in DSB repair post-cleavage that promotes crossover recombination ^11, 12^. Crossover resolution is facilitiated by PRDM9 binding symmetrically to the template (uncut) homolog which generates a NDR within which the DSB-initiating chromosome can stably engage ^13–15^. Despite these observed roles for PRDM9 outside of SPO11 cutting, it is not clear whether PRDM9 influences SPO11 processing or other early steps of meiotic recombination to facilitate synapsis and crossovers. As yeast do not use PRDM9 to establish DSBs, mouse models are necessary to uncover PRDM9 function in infertility and inherited genetic disorders caused by aberrant synapsis and chromosome missegregation.

### END-seq Robustly Detects Mouse Meiotic DSB Hotspots

To probe the early steps of mouse meitotic recombination, we utilized END-seq ^16–18^. In this method, a sequencing adapter is ligated to each end of a DNA break inside an agarose plug after a combination of nucleases ExoVII and ExoT removes ssDNA overhangs. As a result, sequencing reads begin at the terminal end of physiological resection, resulting in libraries of ssDNA-DSB junctions (**Extended Data Fig. 1b**). These exonucleases can be used to detect DSB termini that are either protein-bound or protein-free. For example, etoposide-induced DSBs, which are covalently attached to topoisomerase 2 (TOP2) via an active site tyrosine at the 5’-termini, require ExoVII to remove covalently bound TOP2 ^16^, whereas ExoT can only blunt protein-free overhangs resulting in a ligatable DNA end ^18^. Like TOP2, the topoisomerase-like protein SPO11 remains attached to DSB 5’ ends prior to release by MRE11-mediated nicking as short 20-40 bp oligonucleotides (SPO11-oligos) ^7, 19^ (**Extended Data Fig. 1a**).

We assayed spermatocytes from juvenile mouse testes during the first wave of semi-synchronous meiosis I. We embedded spermatocytes from 20 pooled, 12-14 dpp C57BL/6J (B6) mice in agarose plugs and blunted meiotic ssDNA overhangs with ExoVII and ExoT before ligating sequencing adapters (**Extended Data Fig. 1b**). Approximately 5000 reproducible broken hotspots were called with a threshold of at least 2.5-fold enrichment (**Extended Data Fig. 2a**). We overlapped these END-seq peaks with previously reported B6 meiotic hotspots determined by SPO11-oligo sequencing and DMC1 single-strand DNA sequencing (SSDS) ^8, 11, 20^. By visual inspection and correlative analysis, both END-seq break location and peak intensity overlapped with SPO11-oligos and SSDS, which accounted for 97% and 98% of END-seq peaks respectively (**Fig. 1a, Extended Data Fig. 2b, c**). END-seq therefore provides the first map of directly detected meiotic DSB resection in mammals.

**Figure 1.**
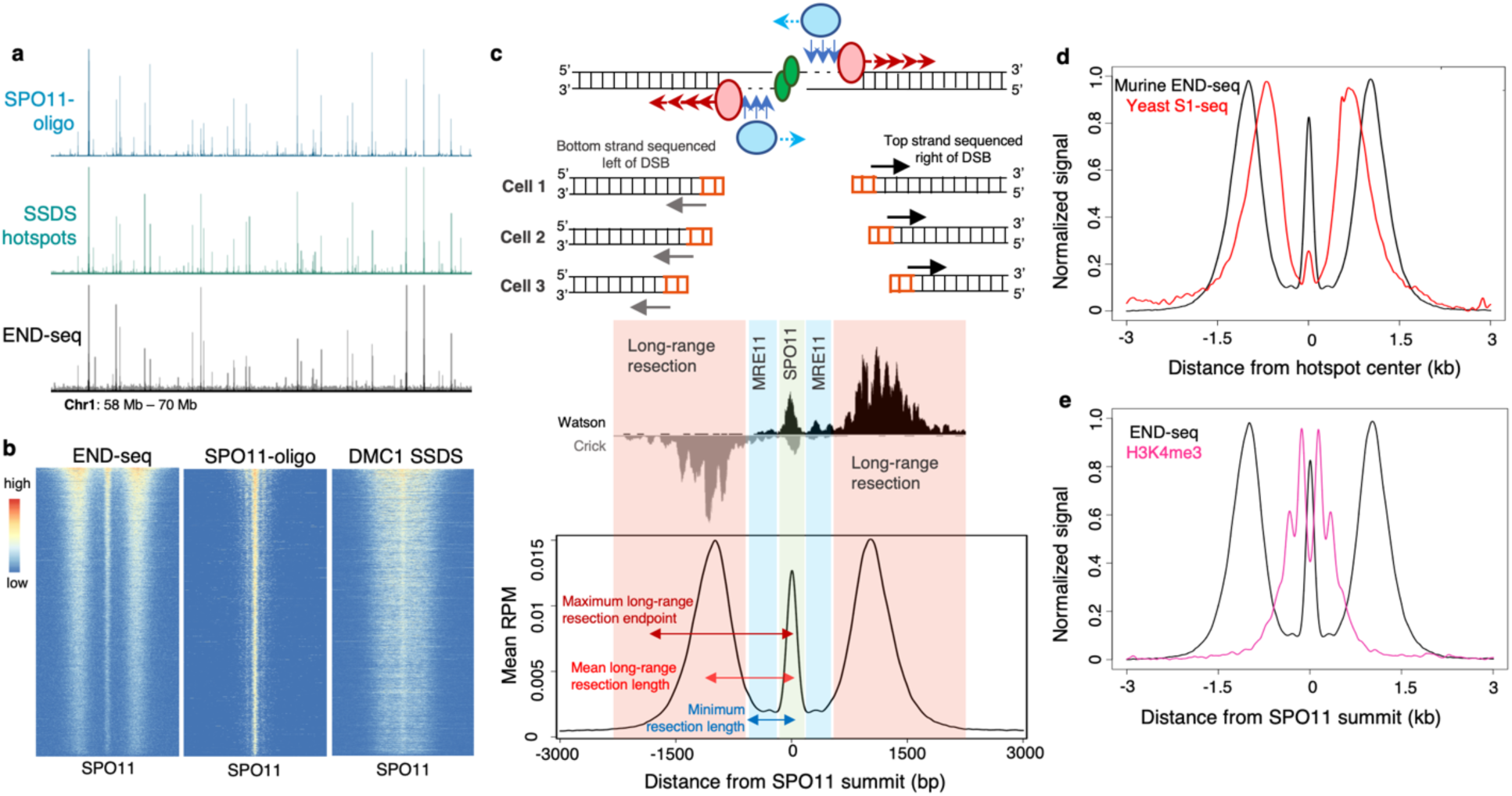
END-seq correlates with previous hotspot mapping techniques and uncovers a uniform pattern of SPO11 processing at all hotspots. (**a**) Representative genome browser profiles of meiotic hotspots on Chromosome 1 for SPO11-oligo sequencing, DMC1 SSDS, and END-seq. Browser axis scales are adjusted to show both hot and cold hotspots simultaneously. (**b**) Heatmaps of END-seq, SPO11-oligo, and SSDS in a ±2.5kb window from hotspot centers (determined by SPO11-oligo summits), ordered by total read count of END-seq, for top 5000 END-seq breaks. (**c**) Schematic representation of the uniform break pattern detected by END-seq, consisting of a central peak at the SPO11 break site, a read-less gap produced by MRE11 mediated-short-range resection and simultaneous engagement of long-range resection, and distal reads indicating the terminal end of long-range resection. The pattern reflects heterogeneity in DSB processing and resection endpoints within the population of spermatocytes (top). This pattern is evident at individual hotspots (middle) and when signal from all hotspots is aggregated (bottom). Minimum resection lengths are calculated by the absence of sequencing reads (blue highlighted region), while mean long-range resection lengths and maximum long-range resection endpoints are calculated by the average and most distal read density from the DSB, respectively (red highlighted region). (**d**) Aggregate signal comparison of murine meiotic END-seq around SPO11-oligo summits and yeast meiotic S1-seq around hotspot centers. (**e**) Aggregated END-seq and H3K4me3 ChIP-seq signal at top 5000 broken WT hotspots (normalized to the same height). Minimum amount of resection (gap region) at all hotspots is mostly confined to H3K4me3.

END-seq captured a strikingly uniform pattern of breakage at all sites that consisted of a strong central peak directly at the site of SPO11 cutting (coincident with mapped SPO11-oligos) with an accumulation of reads flanking the cut site at a defined distance away (**Fig. 1b, c**). We interpret the central peak to be the direct detection of SPO11 breakage while adjacent, distal reads reflect minimum and maximum resection endpoints in the population of spermatocytes (discussed further below).

Using the 2.5-fold enrichment criteria, END-seq peaks called a third of the total hotspots determined by SPO11-oligo seq or SSDS (**Extended Data Fig. 2b, c**). However, END-seq peaks that did not meet this cut-off were nevertheless associated with previously mapped hotspots. For example, 66% of SPO11-oligo loci that were “END-seq negative” nevertheless showed the same DSB detection pattern when these END-seq reads were aggregated (**Extended Data Fig. 3a, b**). Moreover, these END-seq-negative SPO11-oligo sites show a significant reduction in SPO11-oligo reads (**Extended Data Fig. 3c**), indicating that these are the coldest meiotic hotspots. We conclude that END-seq detects breakage at all ∼15,000 previously mapped hotspots, yet the more frequently broken top 33% of hotspots yield the most robust signal. Therefore, unless otherwise stated, the subsequent analyses were performed on the 5,000 strongest END-seq breaks.

### Meiotic DSB Hotspots Are Detectable in a Single Mouse

To determine the sensitivity of the method, we compared 20 pooled juvenile mice to a library made from a single 12 dpp mouse. Both samples called approximately 5000 peaks, with 77-89% shared breaks with highly correlated (r=0.98) END-seq intensities (**Extended Data Fig. 3d**). Importantly, the break pattern at individual hotspots was fully retained in testes from a single mouse (**Extended Data Fig. 3e**). To compare signal-to-noise ratio (S/N) in the two libraries, we followed ENCODE’s assessment of fraction of reads in peaks (FRiP) and cross-correlation profiles (CCPs) for ChIP-seq datasets ^21, 22^. FRiP and CCP values for both libraries exceeded ENCODE’s criteria for signal to noise ratios (**Extended Data Fig. 3f, g**). We conclude that END-seq can accurately assess individual meiotic DSB locations and processing with remarkably little biological material. This high sensitivity bypasses the limitations of other hotspot mapping methods that require either impractical quantities of mice (SPO11-oligo seq) or the availability of species-specific, high-quality antibodies (DMC1 SSDS).

### Estimation of Total Number of DSBs per Cell

To estimate the number of SPO11 DSBs per spermatocyte, we added to each sample a spike-in control consisting of a G1-arrested Ableson-transformed pre-B cell line (*Lig4^−/−^*) carrying a single zinc-finger-induced DSB at the TCRβ enhancer ^16^. This site is expected to break in all spike-in cells, which were mixed in at a 2% frequency with bulk testiscular cells, allowing us to normalize END-seq reads at all ∼15,000 SPO11-oligo hotspots using the signal at the TCRβ enhancer. Based on this approach we calculated approximately fifty total breaks per cell. Since about 15% of 12-14 dpp bulk testicular cells are spermatocytes, as determined by flow cytometry (**Extended Data Fig. 3h**), we estimated that each spermatocyte harbors approximately 350 breaks, consistent with cytological observations of repair protein recruitment to DSB sites ^23^ and genome-wide assessments of cross-over and non crossover events ^15^.

### Increase Breakage in *Atm*^−/−^ Spermatocytes

ATM is thought to negatively regulate SPO11 cutting, and in its absence, hotspot breakage and SPO11-oligos have been shown to dramatically increase in mice ^7^. This ultimately results in early meiotic arrest and apoptosis of *Atm*^−/−^ spermatocytes that carry an excess of unrepaired DSBs ^7^. To further validate this model using direct, genome-wide detection, we performed END-seq on *Atm*^−/−^ spermatocytes that were backcrossed eleven times to the B6 background. Indeed, *Atm*^−/−^ END-seq detected 99% of WT END-seq breaks while calling ∼6.3k new peaks (**Extended Data Fig. 4a**). These novel *Atm*^−/−^ breaks overlapped even better with SPO11-oligos and SSDS sites and showed amplified signal over WT at weak SSDS hotspots (**Extended Data Fig. 4b-d**). These data confirm previous reports that colder hotspots become preferentially hotter in the absence of ATM due to a loss of negative feedback of SPO11 cutting (**Extended Data Fig. 4e**) ^7,8^.

Unexpectedly we found that 7-16% (700 to 1,800) of break sites were not shared in WT SSDS or SPO11-oligo maps respectively (**Extended Data Fig. 4b, c**). Among these were several hundred promoter breaks, typically associated as “default” hotspots in organisms lacking PRDM9 (**Extended Data Fig. 4f**). Indeed, these break locations are among the hottest *Prdm9*^−/−^ sites as determined by SSDS (**Extended Data Fig. 4g**) ^11, 24^. Thus, in the absence of ATM, colder hotspots and default hotspots become increasingly broken. This is consistent with previous observations that ATM-null spermatocytes have increased SPO11-oligo mapping at PRDM9-independent hotspots ^14^.

### The Landscape of Mouse Meiotic DSB Resection

END-seq captures the terminal end of physiologic resection after *in vitro* blunting of the 3’ overhang, with the first nucleotide sequenced corresponding to the position of the ssDNA-DSB junction (**Fig. 1c, Extended Data Fig. 1b**) ^18^. If bidirectional resection of DNA-bound SPO11 by short- and long-range resection machineries are entirely coordinated as in yeast ^4–6^, then the ssDNA-DSB junction will always be beyond the most distal MRE11 nick and correspond to the terminal end of long-range resection (**Fig. 1c, Extended Data Fig. 1b**). In this case, the location of short-range 3’-5’ MRE11 exonuclease activity would not yield any sequencing reads since it does not operate independently of 5’-3’ resection (**Fig. 1c** **and Extended Data Fig. 1b**). Indeed, at every hotspot, we observed a readless “short-range resection gap”, consistent with the tight coupling of resection initiation (by MRE11) and 5’-3’ extension by the long-range resection machinery (**Fig. 1b, c**). The length of this gap reflects the minimum resection endpoints in the spermatocyte population. This corresponds to the maximum distance from the DSB at which MRE11 nicks the strand to be resected plus any constant, minimum distance that 5’-3’ exonucleases traverse. This pattern is reminiscent of S1 nuclease detection of meiotic recombination in yeast ^6^ (**Fig. 1d**). Thus, both SPO11 cutting and its initial processing by coordinated resection mechanisms are highly conserved evolutionary features of meiotic recombination that span unicellular eukaryotes to mammals.

### Nucleosome Positioning Influences Short-Range and Long-Range Resection

The gap size was extremely uniform at all hotspots, with mean maximum distance of 647 nts (**Extended Data Fig. 5a**), and was largely restricted to the two-nucleosome H3K4/K36me3 signal surrounding SPO11 cut sites as determined by ChIP-seq (**Fig. 1e,Extended Data Fig. 5b**). Thus, minimum resection distances correlate well with PRDM9-mediated methylated histone deposition ^10^. Interestingly, the majority of crossover and non-crossover boundaries in mice are also restricted to this region, rarely extending beyond the 650 bp gap (**Extended Data Fig. 5c**) ^15, 25^.

Because minimum resection correlated well with methylated histone deposition, we asked if long-range resection length was similarly determined by nucleosome positioning farther away from SPO11 cutting. Between two replicate END-seq samples, we found high correlation in the pattern of resection among the hotter hotspots detected (**Extended Data Fig. 5d, e**). This strong reproducibility prompted us to assess the distribution of resection subpeaks within hotspots. These breaks exhibited distinct subpeaks within long-range resection signal and were distributed apart from each other at a median distance of 210 nucleotides (**Extended Data Fig. 5f**). This suggests that nucleosome positioning also influences the endpoints of long-range resection, consistent with findings in budding yeast ^6^.

### Minimal Roles of EXO1, BRCA1, and 53BP1 in Long-Range Resection

Long-range resection end points showed greater variation (1-3 kb) relative to the minimum resection gap region (0.4-1kb) (**Extended Data Fig. 5a**). Previous studies estimated total mammalian meiotic resection lengths based on the extent of DMC1 bound to ssDNA overhangs as measured by DMC1 SSDS ^8^. Strikingly, we found that maximum resection end points extended significantly farther at all hotspots than DMC1-bound ssDNA (**Fig. 2a, b**). ssDNA occupied by DMC1 ranged from 800-2700 nts, whereas maximum long-range resection lengths (defined in **Fig. 1c**) determined by END-seq were 1.2-1.6 fold greater (**Fig. 2b, Extended Data Fig. 6a**). These data indicate that DMC1 binds to only a portion of the available ssDNA and underestimates the total extent of meiotic resection. One potential explanation is that single tracks of ssDNA can be co-occupied by DMC1 and RAD51 filaments with DMC1 loaded more proximally to the break site than RAD51 ^26^. While unlikely, this difference may also be due to technical limitations in SSDS library preparation that relies on microhomology-mediated hairpin formations naturally present in ssDNA tracks ^27^.

**Figure 2.**
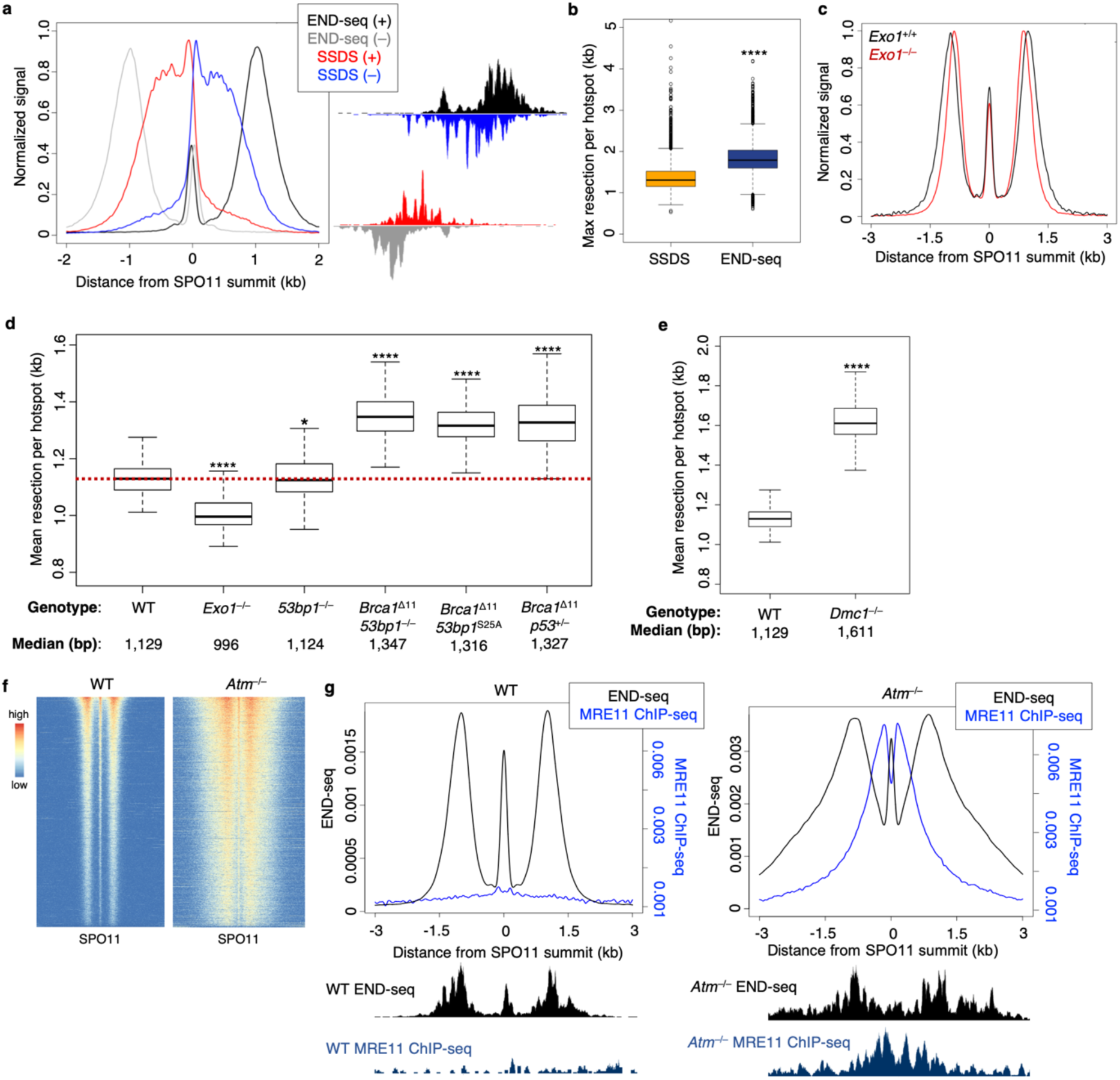
END-seq accurately measures hotspot resection lengths and elucidates resection regulation. (**a**) Resection length comparison between END-seq and DMC1 SSDS. Top (+) and bottom (–) strand distributions of END-seq and SSDS coverage show increased resection detection by END-seq (left, signal normalized to the same height) and is evident at individual hotspots (right). (**b**) END-seq detects increased maximum long-range resection compared to SSDS genome-wide. *p < 0.01; ****p < 1e-10; T test. (**c**) Aggregate plot of END-seq signal in WT and *Exo1*^−/−^ spermatocytes (signal normalized to the same height). (**d**) Comparison of mean long-range resection between WT, *Exo1*^−/−^, *53bp1*^−/−^, *Brca1^Δ11^53bp1*^−/−^, *Brca1^△11^53bp1^S25A^*, and *Brca1^△11^p53^+/–^* at top 250 hotspots. *p < 0.01; ****p < 1e-10; T test. (**e**) Comparison of mean long-range resection between WT and *Dmc1*^−/−^ at top 250 hotspots. ****p < 1e-10; T test. (**f**) Heatmaps of WT and *Atm*^−/−^ spike-in normalized END-seq ±5kb around SPO11-oligo summits for top 5000 hotspots. (**g**) Hotspot aggregated non-normalized signal for END-seq and MRE11 ChIP-seq in WT (left) and *Atm^−/−^* (right). Individual hotspot examples are shown below.

Because the resection pattern in mouse mirrored so closely that observed in yeast (**Fig. 1d**) ^6^, we wanted to know whether they utilized the same long-range end processing machineries. *Exo1*-deficient yeast exhibit total loss of 5’-3’ long-range resection ^3^. In contrast, EXO1 and DNA2 act redundantly in yeast vegetative and mammalian somatic cells to mediate end resection ^28–30^. Surprisingly, END-seq analysis of juvenile *Exo1*^−/−^ spermatocytes revealed that long-range resection was largely intact (**Fig. 2c**). When averaged genome-wide, we found a small, though significant, reduction in long-range resection distance in *Exo1*^−/−^ cells (median resection tract: WT, 1,129 nts, vs. *Exo1*^−/−^, 996 nts (**Fig. 2d**). We therefore conclude that compared to yeast, mammalian meiosis has evolved additional mechanisms to achieve extensive 3’ overhangs, perhaps through utilization of redundant DNA2 exonuclease activity ^28–30^.

In somatic interphase cells, 53BP1 has been shown to inhibit long-range resection of DSBs ^31^. One principle role of BRCA1 is to counteract 53BP1’s block to resection in S phase, possibly by excluding it from chromatin proximal to DNA damage sites ^32^. In addition, BRCA1 acts post-resection to load the RAD51 recombinase onto 3’ ssDNA ^33^. Despite these extensive studies demonstrating their importance in somatic cells, the roles of 53BP1 and BRCA1 in regulating meiotic resection are unknown. We therefore performed END-seq on *53bp1*^−/−^ and BRCA1-deficient spermatocytes and measured overall resection lengths. In striking contrast to 53BP1-deficient somatic cells in which DSB resection is consistently and acutely increased ^18, 31, 34^, we found that resection lengths were comparable at all meiotic hotspots in *53bp1*^−/−^ and WT spermatocytes (**Fig. 2d**).

We then assayed *Brca1^Δ11^p53*^+/–^ mice, which exhibit known defects in BRCA1 function yet are alive due to partial p53 apoptotic suppression ^35^. Contrary to our expectations ^31, 36^, *Brca1^Δ11^p53*^+/–^ spermatocytes showed a mild increase in DSB resection relative to WT controls (**Fig. 2d**). Moreover, *Brca1^Δ11^53bp1^−/−^* and *Brca1^Δ11^53bp1^S25A^* ^36^ spermatocytes exhibited a similar increase in resection as *Brca1^Δ11^p53*^+/-^ spermatocytes (**Fig. 2d**). Based on these findings, we conclude that in contrast to their well-defined antagonistic roles in DSB processing in interphase cells, 53BP1 does not inhibit resection while BRCA1 does not promote resection during meiotic recombination. As discussed below, one reason why end resection increases (rather than decreases) in BRCA1-deficient cells may be due to inefficient loading of the recombinase. Consistent with this, *Brca1^Δ11^p53*^+/–^ spermatocytes have reduced RAD51 and DMC1 focus counts ^37^.

### RAD51, DMC1, and ATM Negatively Regulate Long-Range Resection of DSBs

In budding yeast, cells lacking DMC1 show hyperresection, perhaps due to a negative feedback on resection mediated by recombinase loading ^4, 6^. Given our results that BRCA1-deficient spermatocytes exhibited increased resection (**Fig. 2d**) associated with defective RAD51/DMC1 foci ^37^, we hypothesized that RAD51 or DMC1 loading onto ssDNA might limit long-range resection. Consistent with this, *Dmc1*^−/−^ spermatocytes showed substantially increased resection relative to WT at all hotspots (**Fig. 2e, Extended Data Fig. 6b, c**). Minimum resection lengths also increased in *Dmc1*^−/−^ cells by approximately 400 nts relative to WT (**Extended Data Fig. 6b, c**). Since it is unlikely that MRE11 endonuclease activity, the earliest step in SPO11 processing ^2^, is confined by DMC1, we interpret this to mean that DMC1 loading post-resection limits both the minimum and maximum length that the long-range resection machinery processes DSBs. DMC1/RAD51 loading onto ssDNA might limit resection by preventing the re-initiation of exonucleases on already resected ssDNA.

ATM negatively regulates DSB induction by SPO11 ^7^. Despite the fact that *Atm*^−/−^ spermatocytes harbor a 10-fold increase in DSBs ^7^, RAD51/DMC1 foci form at similar levels as in WT while RPA foci counts are increased in mutant cells ^38^. One possible reason why the number of foci do not correlate with the large increase in DSBs could be that recombinase levels or activity is limiting for filament formation. In this scenario, when excess DSBs are generated in an ATM-null background, the pool of RAD51 and/or DMC1 is insufficient to load onto all ssDNA regions. Inefficient loading would result in a concomitant increase in resection lengths if recombinase filament formation negatively regulates resection. In accord with this hypothesis, the mean maximum resection lengths increased almost 2-fold in *Atm*^−/−^ spermatocytes relative to WT (WT: 1,845 nts. vs. ATM: 3,351 nts; **Fig. 2f, Extended Data Fig. 6d**). Moreover, in ATM knockouts, resection lengths were significantly greater and more widely distributed than even DMC1 knockouts (**Extended Data Fig. 6c, d**). This suggests the possibility that the increase resection associated with ATM kinase deficiency may not only be due to recombinase “exhaustion,” but also because of the additional influence of nucleosome positioning on resection (**Extended Data Fig. 5d-f**).

### ATM Coordinates Short- and Long-Range Resection

In addition to the hyper-resection observed at a subset of breaks, distinct boundaries between the short- and long-range resection practically disappeared in *Atm*^−/−^ spermatocytes, resulting in reads mapping within the gap region (**Fig. 2f, Extended Data Fig. 7a**). The decreased minimum resection could indicate that ATM regulates the tightly coupled activities of short-range and long-range resection, which in WT generates ssDNA-DSB junctions beyond the most distal MRE11 nick (**Extended Data Fig. 7b**). This could reflect a role for ATM in promoting normal MRE11 resection initiation perhaps through CtIP phosphorylation, as suggested previously for Tel1-mediated Sae2 phosphorylation in yeast ^6^. In either case, DSB processing would be incomplete in the absence of ATM.

If some DSBs are not resected in ATM-deficient spermatocytes, we would predict that MRE11 would accumulate at unprocessed DNA ends. Consistent with this hypothesis, ChIP-seq for MRE11 revealed significant chromatin binding at all hotspots in *Atm^−/−^* cells (**Fig. 2g**, **right panel, Extended Data Fig. 7c**). In contrast, we detected no signal in similarly broken regions in WT cells (**Fig. 2g, left panel**), indicative of completed resection. Strikingly, MRE11 signal was predominantly within the END-seq gap and greatly diminished in the H3K4me3 nucleosome-depleted region (NDR) in *Atm^−/−^* cells (**Extended Data Fig. 7c**). As expected, its binding also correlated with END-seq intensity (**Extended Data Fig. 7d**). This supports the idea that ATM promotes MRE11 activity, which in turn coordinates the processing of DNA bound SPO11. Notably, the recruitment of MRE11 to breaks in ATM-null cells is localized predominantly to the WT read-less gap occupied by H3K4me3/K36me3 (**Fig. 1e, Extended Data Fig 7e**). This further supports the idea that the gap reflects MRE11-dependent end-processing.

### END-seq Central Peak Reflects DNA-Bound SPO11

At all hotspots, END-seq detected a uniform accumulation of reads aligned to the center of the DSB. This central peak was coincident with mapped SPO11-oligos, having a width (400 bp) similar to SPO11-oligo seq hotspots (300-400 bp) and was restricted to the NDR of H3K4/K36me3 (**Fig. 3a, 1e, Extended Data Fig. 5b**) ^8^. Moreover, even low-level secondary oligos that are adjacent to the central SPO11 hotspot peak ^8^ were detectable by END-seq (**Fig. 3a, right**). This suggested that the central peak represented a fraction of the total DSBs in the spermatocyte population in which SPO11 is not yet released, thereby highlighting the heterogeneity of DSB processing.

**Figure 3.**
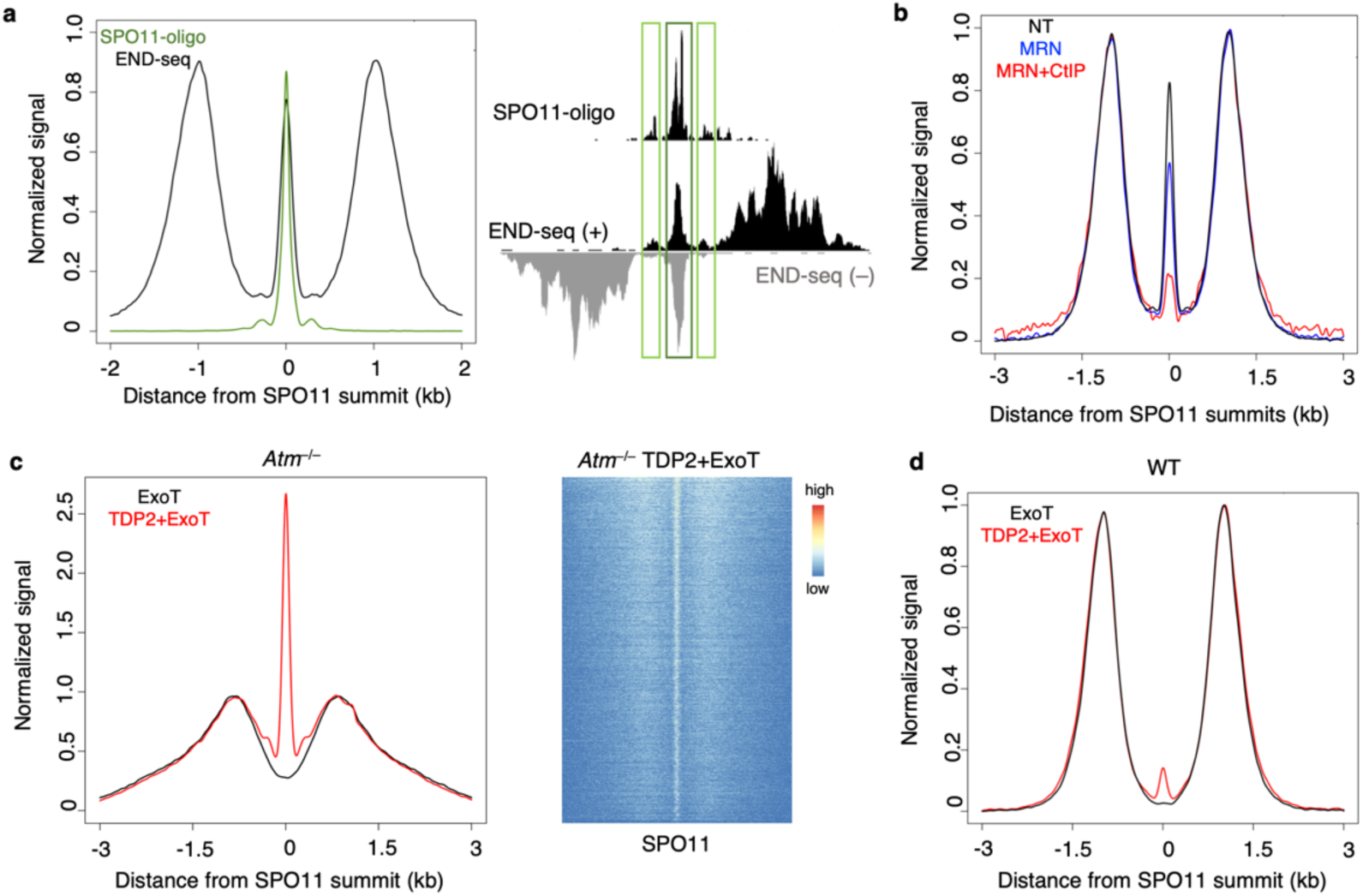
WT and *Atm^−/−^* hotspots contain distinct species of chromatin-bound SPO11. (**a**) END-seq central peak and SPO11-oligo reads are coincident as evidenced by aggregated signal around SPO11-oligo summits (left, normalized to the same height) and at individual hotspots (right). Both the primary SPO11 peak and the adjacent secondary peaks are highlighted. (**b**) Pretreatment with purified human MRN+CtIP reduces ExoVII+ExoT central peak detection (red line) over no pretreatment (NT, black line) and depends on the presence of CtIP (blue line). (**c**) END-seq with ExoT blunting alone also shows no central peak in *Atm*^−/−^ cells (black line). Pretreatment with purified human TDP2 before ExoT processing massively recovers SPO11 signal, indicating abundance of unresected, bona fide SPO11 cleavage complexes (red line). Aggregate plot (left, normalized to same resection height) and heatmap (right) in a ±3kb window around SPO11-oligo summits. (**d**) END-seq aggregate plot of WT ExoT blunting with (red line) and without (black line) TDP2 pretreatment shows only minor recovery of SPO11 signal, suggesting that WT SPO11-bound DSBs are not simply unresected cleavage complexes.

If 5’ covalently-bound SPO11 DSBs existed at the time of END-seq processing, then the proteinase K digestion would leave behind a two-bp 5’ overhang with a phosphotyrosyl bond that requires ExoVII digestion to fully blunt the end, analogous to our studies on TOP2 cleavage complexes (TOP2cc) ^16^. To test this hypothesis, we performed END-seq with ExoT blunting only, allowing adapter ligation only to protein-free DNA ends while excluding ends with protein adducts, such as any remaining SPO11cc. Strikingly, ExoT detected only fully resected DNA ends with a total absence of central signal at all hotspots (**Extended Data Fig. 8a, b**). These data indicate that the central peak, which represents approximately 11% of the total DSB signal and is present all hotspots, reflects SPO11 covalently bound to its break site.

To validate that the ExoVII-dependent central peak was due to SPO11 bound to the break and not some other form of occlusion, we speculated that it might be possible to remove SPO11 tyrosyl-linked DNA through incubation with purified human MRE11/RAD50/NBS1 (MRN) and CtIP prior to blunting with ExoVII+ExoT, mimicking the *in vivo* processing of meiotic DSBs. Indeed, we observed a dramatic loss of central signal by preincubation with MRN and CtIP (**Fig. 3b**) ^39^. Incubation with MRN alone (prior to ExoVII+ExoT) did not efficiently remove the central peak (**Fig. 3b),** consistent with the finding that CtIP is required for MRE11 endonuclease processing of protein-bound DSBs ^40, 41^. Thus, purified MRN/CtIP recognizes and removes the remaining SPO11 phosphotyrosyl bonds associated with the central signal. These data therefore support the idea that a fraction of SPO11 remains physiologically bound to a subset of breaks (11%) at virtually all hotspots after cutting.

### Increased Fraction of Unresected SPO11-Bound DSBs in *Atm*^−/−^ Spermatocytes

Because Tel1 regulates MRE11 initiated resection, there is a dramatic increase in unresected DSBs in *TEL1*-deficient yeast ^6^. If ATM functions similarly to Tel1, then we would expect an accumulation of SPO11cc, well above WT levels, at the center of the hotspots. The abundant MRE11 ChIP-seq signal specifically in *Atm***^−/−^** cells (**Fig. 2g**), indicative of incomplete processing, would also predict a vast increase in SPO11cc in the mutants; however, we observed that the central peak detected by ExoVII+ExoT was similar in WT and ATM-null cells (**Extended Data Fig. 7a**).

We therefore sought to modify END-seq to specifically probe for SPO11cc. We hypothesized that preincubation with purified tyrosyl-DNA phosphodiesterase 2 (TDP2) ^42^ would remove the remaining phosphotyrosyl adduct after proteinase K treatment, generating a two-nucleotide, protein-free 5’ overhang that ExoT could readily blunt for adapter ligation. While ExoT alone detected no central peak in ATM-null cells, similar to WT (**Fig. 3c, Extended Data Fig 8a**), TDP2+ExoT END-seq captured an astonishingly robust central signal (**Fig. 3c**). Moreover, this peak was strongly detected at all hotspots (**Fig. 3c, right**). SPO11cc detection by TDP2+ExoT far exceeded the efficiency of detection with ExoVII+ExoT both at autosomes and at the non-PAR X chromosome (**Extended Data Fig. 8c, d**), most likely reflecting the biochemical preference for TDP2 over ExoVII to remove the phosphotyrosyl bonds and allow for adapter ligation ^42^. These data indicate that ATM-deficient spermatocytes accumulate unresected SPO11cc, similar to yeast lacking Tel1 ^8^.

In WT cells, the ExoVII+ExoT central signal, which is also SPO11-bound, represents 11% of the total DSB fraction (**Fig. 3a**). Yet, combination of TDP2 and ExoT only recovered 16% of the ExoVII+ExoT detected fraction of central signal found in WT cells (**Fig. 3d**), corresponding to 0.5-2% of the total DSB signal. Given the high efficiency of TDP2+ExoT in ATM-null cells, it was initially unclear why the WT TDP2+ExoT central signal was so low. We speculated that in WT cells, the central signal might not simply reflect unresected SPO11cc, as observed in *Atm*^−/−^ spermatocytes. Rather, there might be an additional structure at the break site associated with SPO11-bound DNA in WT cells that somehow prevented recognition by TDP2.

### DNA-Bound SPO11 is Dependent on Homolog Engagement and PRDM9

Our first clue to understanding the distinct nature of SPO11-bound DNA in WT *vs. Atm^−/−^* cells was the observation that it was missing from all hotspots in *Dmc1^−/−^* spermatocytes (**Fig. 4a**). This prompted us to ask whether the central peak associated with DNA-bound SPO11 was also dependent on ssDNA strand invasion into the homologous partner. Remarkably, we found a loss of central signal at all non-PAR X chromosome hotspots (**Fig. 4b**), which repair from the sister chromatid since the X chromosome has no homolog in males. The residual central signal resembled the TDP2+ExoT signal in WT cells (**Fig. 3d**) and therefore might simply reflect the amount of unresected SPO11 naturally present on all chromosomes. In striking contrast, the central signal associated with autosomes and the non-PAR X chromosome was virtually identical in *Atm^−/−^* spermatocytes, again confirming that the central signal is largely SPO11cc in the absence of ATM (**Extended Data Fig. 8e**).

**Figure 4.**
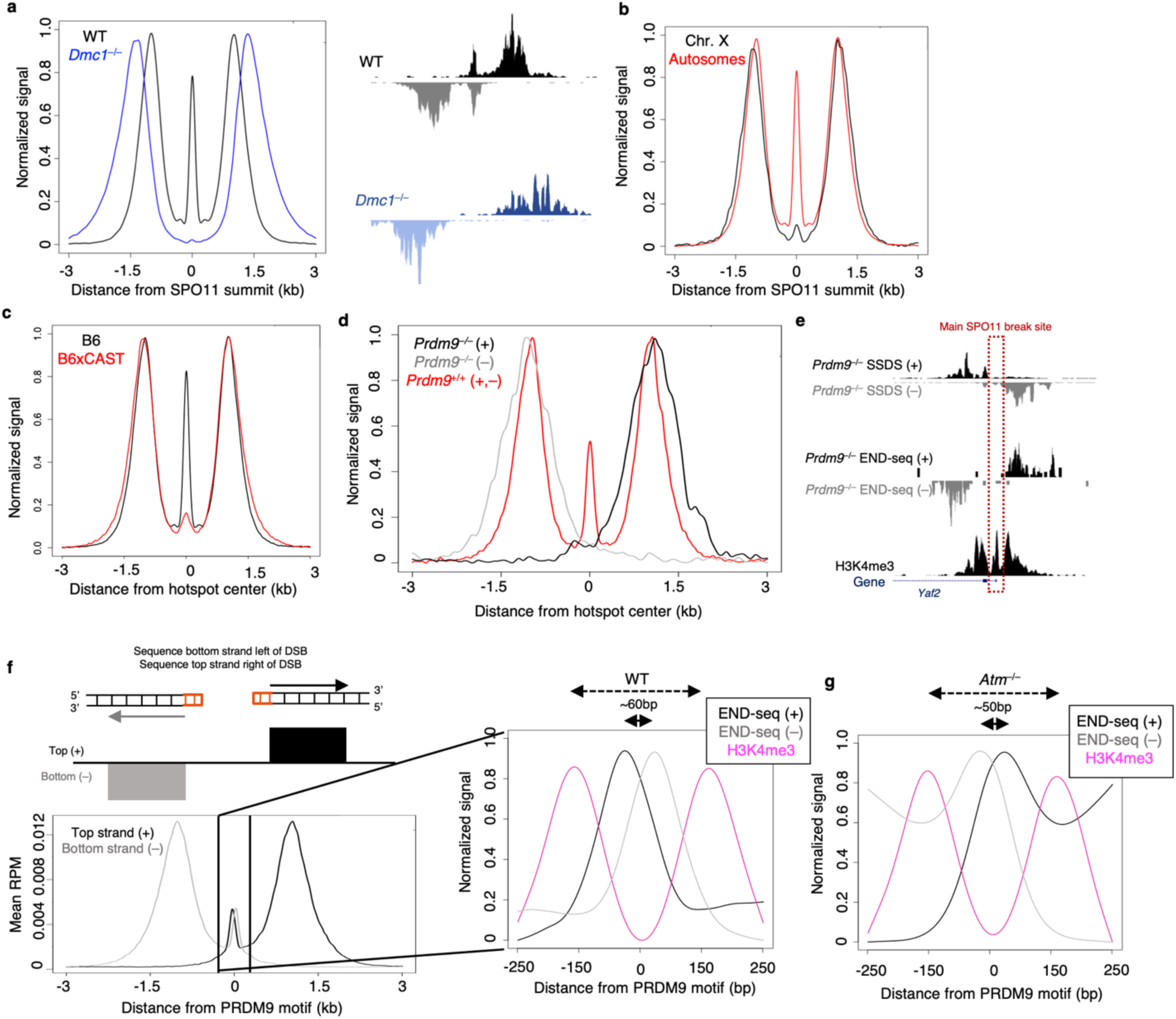
WT central peak detection relies on homolog engagement and PRDM9. (**a**) Aggregated signal (left, normalized to the same height) and hotspot example (right) showing absence of SPO11 central peak in *Dmc1*^−/−^. (**b**) Reduction in central signal at non-PAR X chromosome hotspots; aggregated signal on ChrX compared to all autosomes. (**c**) Aggregated signal of B6 (centered on B6 SPO11-oligos) versus B6xCAST hybrid (centered on hybrid SSDS hotspot centers). (**d**) SPO11 central peak is absent in *Prdm9^−/−^* END-seq signal aggregated around default SSDS hotspot centers at ∼200 sites with least overlap in SSDS top and bottom strands. WT END-seq is centered around WT SPO11-oligos at PRDM9-dependent hotspots. (**e**) *Prdm9^−/−^* SSDS and END-seq tracks at a single default hotspot with minimal SSDS top and bottom strand overlap. Main SPO11 break site (red dotted line) is inferred from SSDS pattern. (**f**) At fully processed and resected SPO11 DSBs, END-seq top and bottom strand reads exhibit a “correct” polarity to the right and left of the DSB, respectively (left). WT reads at the center of hotspots that originate from a subset of DSBs with bound SPO11 show a top and bottom strand polarity that is reversed from what is expected (right, zoomed in at NDR). (**g**) *Atm^−/−^* END-seq central peak has correct polarity of top and bottom strands within the NDR. Strands show significant separation of ∼50 bp, likely indicating multiple SPO11 cuts within the same hotspot.

The analysis of hybrid mouse strains with different PRDM9 alleles has revealed that the degree of asymmetry in PRDM9 binding—that is, whether PRDM9 binds unequally to both homologs-predicts increased asynapsis and hybrid infertility ^13, 14^. When PRDM9 fails to bind the unbroken homologous chromatid, there is a severe reduction in both crossover and noncrossover events ^14, 15^. Indeed, it has been suggested that similar to the X chromosome, asymmetric hotspots might be repaired from the sister chromatid ^15^. We therefore performed END-seq on spermatocytes derived from juvenile B6xCAST hybrids with divergent genomes. Strikingly, the central peak was reduced to 2-3% of the total DSB signal in B6xCAST compared to the 11% observed in B6 (**Fig. 4c**) and much lower than in CAST alone (**Extended Data Fig. 8f**). Based on these findings, we conclude that the central signal detected by ExoVII+ExoT in WT spermacotyes is associated with DNA-bound SPO11, and is dependent on the degree of homologous chromosome engagement.

In mice lacking PRDM9, DSBs occur at H3K4me3 sites mainly associated with promoters ^9, 11, 12^. However, these DSBs are not repaired efficiently as crossovers resulting in meiotic arrest ^11^. To test how PRDM9 deficiency impacts homolog engagement and resection we performed END-seq on *Prdm9*^−/−^ spermatocytes. As SPO11 can generate multiple breaks within the H3K4me3 sites at promoters, we focused our analyses on *Prdm9*^−/−^ SSDS sites that exhibited the least overlap between top and bottom strand reads, i.e. hotspots that are most likely to have one main SPO11 cut site within the promoter (**Extended Data Fig. 9a**). Such SSDS hotspots showed limited signal at the center, in line with these sites having a strong preference for a single SPO11 cut site (**Extended Data Fig. 9a, b**). Strikingly, END-seq analysis of *Prdm9*^−/−^ spermatocytes revealed a total absence of the central peak at hotspots with the least SSDS strand overlap, examined either on aggregate or individually in the genome browser, whereas short-and long-range resection appeared to be relatively intact (**Figures 4d, e, Extended Data Fig. 9c**). The absence of central signal is consistent with the idea that PRDM9 promotes homolog engagement, which in turn facilitates crossovers.

### DNA-Bound SPO11 is Associated with Recombination Intermediates in WT Cells

At a meiotic DSB that has been fully resected on both sides and SPO11 completely released, adapters ligated to the right end of the break will align to the top (+) DNA strand, whereas adapters ligated to the left end will align to the bottom (–) strand (**Fig. 1c;** **Fig. 4f, left**). When reads from all hotspots are aggregated, the resection signal around SPO11 cuts exhibited this “correct” polarity for the top and bottom strands (**Fig. 4f, left**). However, a close examination of the WT END-seq reads associated with the central signal unexpectedly revealed a “wrong” polarity, in which top strand reads aligned slightly to the left within the NDR and bottom strand reads aligned slightly to the right (**Fig. 4f, right**). As SPO11 generates a DSB with only 2 nt overhangs, if the central signal were merely a collection of unresected SPO11cc, then aggregating the top and bottom strand DSB endpoints should show no separation between them. However, we observed a top and bottom strand shift of ∼60 nts in the “wrong” orientation of what is expected for DSBs (**Fig. 4f, right**). Detecting a signficant reverse in the expected polarity indicated that the central peak was not simply SPO11cc. This suggested that while SPO11 remains bound to a fraction DSBs, there is some kind of asymmetry associated with DNA-bound SPO11 that influenced END-seq detection compared to when SPO11 is released and both DNA ends are fully resected (**Fig. 4f**).

One potential mechanism that could contribute to the wrong polarity is asymmetric processing of SPO11. That is, if the two DNA ends bound by SPO11 are processed at different efficiencies by MRE11, one end might be incompletely processed, leaving SPO11 covalently bound to its cut site, while the other end is processed to completion and SPO11-oligo released. In this scenario, only the incompletely processed SPO11-bound end would contribute to central peak signal. The fully processed end (**Extended Data Fig. 10a,** top, left end of the DSB) would result in the release of the SPO11-oligo that would generate a protein-free 3’ overhang. This (SPO11-free) DNA end would in turn be blunted by END-seq and sequencing reads would be detected within the distal, long-range resection peaks (**Extended Data Fig. 10a**, bottom, left end of the DSB). In contrast, the other side of the DSB would be incompletely resected by MRE11 and retain SPO11-covalently bound to a two-nucleotide, 5’ overhang (**Extended Data Fig. 10a,** top, right end of the DSB). END-seq detection (with ExoVII+ExoT) would then remove SPO11 and sequence the remaining dsDNA, with the first nucleotide sequenced being the SPO11 break site itself (**Extended Data Fig. 10a,** bottom, right end of the DSB). This would result in top and bottom strand central peak reads with reversed polarity within the NDR, as SPO11 breaks to the left of the NDR would contribute top strand reads aligning left of center, and SPO11 breaks to the right would contribute bottom strand reads aligning right of center (**Extended Data Fig. 10a**). In a population of spermatoctyes in which there is cutting on both sides of the hotspot center, END-seq would detect an overall signal of resection reads with correct polarity and central reads with reversed polarity (**Extended Data Fig. 10a,** bottom).

How could such asymmetric MRE11-mediated processing arise? Most SPO11-oligo sequencing reads cluster in the center of the nucleosome free depleted region where PRDM9 is also bound, suggesting that PRDM9 does not block SPO11 access ^8, 43^. Rather, we imagine that DNA-bound PRDM9 may guide the position at which SPO11 cuts within the nucleosome-free region, which might be slightly displaced on average by 30 base-pairs (half the size of 60 bp shift) from PRDM9 itself (**Extended Data Fig. 10a**). It has been suggested that PRDM9 often remains bound on the uncut chromosome ^14^ while SPO11 has been proposed to be associated with DNA ends ^19^ until or even subsequent to strand invasion ^14, 19^. If PRDM9 similarly remains bound post-cleavage between the SPO11 cut and MRE11-endonucleolytic nicking position, it could interfere with MRE11 release of SPO11 via 3’-5’ resection. This would prevent MRE11 from generating a fully ssDNA overhang only on one side of the DSB (**Extended Data Fig 10a,** top). Because the natural on/off binding affinity of PRDM9 would determine the frequency at which MRE11 activity is blocked and SPO11-bound DNA is captured by END-seq, we would expect that the central peak to be detected at all hotspots genome-wide, as observed (**Fig. 1b**). Moreover, all hotspots had equal ratios of central peak to resection signal (∼11%), indicating that no hotspot had preference over others, regardless of break frequency.

The central signal not only reflected asymmetric MRE11-mediated processing, but also required DMC1-mediated strand invasion and engagement with the homologous chromosome template (as shown above). Due to this dependency, we infer that SPO11 remains bound post homolog engagement and during the formation of a recombination intermediate (RI). We therefore refer to this RI, with SPO11 capped to the 3’ resected end, as SPO11-RI (**Extended Data Fig. 10a**).

### Increased SPO11 double-cutting at the same hotspot in *Atm*^−/−^ spermatocytes

The central signal in ATM-null spermatocytes is largely comprised of unresected SPO11 (SPO11cc), which accumulates MRE11 *in vivo* and is sensitive to TDP2-mediated processing (**Fig. 2g,** **Fig. 3c, Extended Data Fig. 8c, d**). This is distinct from the enrichment of SPO11-RI observed in WT cells. Elevated levels of SPO11cc could arise from decreased MRE11 endonucleolytic or 3’-5’ exonuclease activity (**Fig. 2f, g; Extended Data Fig 7**), which is also characteristic of yeast *Tel1* deficiency ^6^. If SPO11cc reflects fully unresected DSBs, the END-seq signal should have the correct polarity, similar to resection endpoints, because the top and bottom strands, still bound by SPO11, would be equally susceptible to ExoVII-mediated processing. To examine this, we strand separated the central signal in *Atm*^−/−^ cells. As predicted, we observed the correct polarity expected for canonical DSBs (**Fig. 4g**, bottom strand, left; top strand, right). Unexpectedly, the strands still exhibited a 50 base-pair gap within the nucleosome-depleted region (**Fig. 4g**). If SPO11 cut once on each chromatid throughout the NDR within the population of cells and remained bound to DNA, there would be a 2-bp gap between top and bottom strand DSB endpoints. We therefore infer that the larger gap size reflects SPO11 double-cutting within the same hotspot (**Fig. 4g, Extended Data Fig. 10b**) ^44^. These measurements are consistent with the increased 40-70 nucleotide SPO11-oligo species that were detected in ATM-null mice ^7^, as two distinct SPO11 cuts adjacent to one another could release these longer oligos without MRE11 endonuclease activity (**Extended Data Fig. 10b**). Moreover, MRE11 ChIP-seq revealed a notable dip in MRE11-binding exactly within the NDR (**Fig. 2g, Extended Data Fig 7c**), consistent with the loss of DNA within hotspot centers. These results are supported by genetic evidence of double-cutting in *Atm^−/−^* spermatocytes (A. Lukaszewicz and M. Jasin, personal communication).

Increased double-cutting around PRDM9 binding sites would preclude it from blocking any MRE11 short-range resection that does occur, thereby reducing the frequency of SPO11-RIs in ATM-null cells (**Extended Data Fig. 10b**). Our finding that central signal in *Atm^−/−^* spermatocytes exhibits the correct polarity (**Fig. 4g**), and that this signal is identical on the autosomes and non-PAR X chromosome (**Extended Data Fig. 8e**), is consistent with a significant reduction in SPO11-RI. Therefore, through its regulation of SPO11 cutting and resection, ATM indirectly regulates the formation of SPO11-RI.

## Discussion

We show here the capacity of END-seq to elucidate early meiotic pathways that are critical for proper chromosome segregation and fertility. Because the method requires little starting material (as low as a single mouse) and can be modified through differential enzymatic reactions to detect distinct DSB structures, we are able to bypass the limitations imposed by previous meiotic hotspot profiling techniques. In doing so, we uncovered a pattern of resection, strikingly uniform at all DSBs. This reflects the fact that short- and long-range resection are tightly coupled in a single processive reaction that is mediated by ATM. This pattern is highly reminiscent of meiotic resection in yeast, and reflects the evolutionary conservation of DSB processing pathways.

While yeast do not possess obvious homologs of BRCA1, BRCA1 has a well-described function in supporting the resection of DSBs in somatic mammalian cells ^33^. Surprisingly, we find that BRCA1 does not similarly promote resection during meiotic recombination. One potential reason could be that 53BP1 is not recruited to DSBs in mitotic cells or in early prophase meiocytes ^45, 46^, which may reflect similarities in pathways that suppress DSB repair during mitosis and meiosis. If the primary function of BRCA1 is to counteract 53BP1’s block to resection at DSBs, this function of BRCA1 would not be needed during mitosis or meiosis.

In addition to promoting resection in somatic cells, BRCA1 also facilitates the loading of RAD51/DMC1 onto ssDNA ^47^, a function which appears to be conserved in meiosis ^37^. In several BRCA1-deficient mouse strains with known defects in RAD51/DMC1 filament formation, we observed a slight increase in resection tracts. We suggest that this reflects an indirect role for BRCA1 in limiting resection by promoting recombinase loading onto ssDNA. Consistent with this idea, *Dmc1*^−/−^ spermatocytes displayed a dramatic increase in minimum and maximum resection endpoints. We additionally observe hyper-resection in ATM-null spermatocytes, which we propose is due in part to the limited availability of recombinases to form filaments in the prescence of excessive SPO11 cutting. Thus, we imagine that in BRCA1-, DMC1-, and ATM-deficient spermatocytes, defective recombinase loading permits reiterative engagement of the long-range resection machinery.

While some aspects of resection are highly conserved in yeast and mammals, SPO11-processing appears not to be. Unlike yeast, mammalian cells accumulate significant levels of DNA-bound SPO11 that represent both unresected SPO11 cleavage complexes (SPO11cc) and a recombination intermediate with 5’ covalently bound SPO11 (SPO11-RI). Our model suggests that incomplete processing of SPO11 in mammals, perhaps due to PRDM9 blocking, still allows for, and may even promote, strand invasion via DMC1 into a homologous template (**Extended Data Fig. 10a**). Such a structure, with SPO11 bound to a small stretch of dsDNA, would grant stability to the large ssDNA tract generated, making it a suitable substrate for end-ligation and sequencing. While SPO11-RI is readily detectable by END-seq using ExoVII+ExoT and represents 11% of the total DSB signal, processing with TDP2+ExoT only detects SPO11cc (**Fig. 3c, d**). We therefore estimate that out of the total DSB signal detected by ExoVII+ExoT in WT cells, 0.5-2% is truly unresected SPO11cc and the remaining ∼10% is SPO11-RI.

What is the biological relevance of SPO11-RI? One possibility is that SPO11-RI facilitates non-crossover (NCO) and/or cross-over (CO) events necessary for the exchange of genetic information between homologs. The majority of NCO and CO events occur only in the 4,000 hottest hotspots ^15^, and we have found that SPO11-RI occurs across all 5,000 DSBs detected by END-seq (**Fig. 1b**). NCO and CO events are reduced in strains with asymmetric PRDM9 hotspots, and we found strong depletion of the central signal in hybrid strains. CO are less likely to occur in gene rich regions ^15^, and we have found that the central signal is decreased with increasing gene expression independent of DSB formation (**Extended Data Fig. 10c, d**). Finally, the positioning of NCO and CO tracts are highly enriched in a small region (generally <500 bp) surrounding the PRDM9 motifs ^15, 25^. It is possible that the “capping” of the 3’ ssDNA by SPO11 (**Extended Data Fig 10a**) prevents further polymerization or double Holliday junction migration, therefore restricting SPO11-RI to a 400 bp region surrounding the PRDM9 motif. Although SPO11-RI correlates with NCO and CO events, additional studies will be necessary to determine its precise physiological function.

The loss of the central signal in mouse hybrids, in *Dmc1^−/−^* mice, and on the WT non-PAR X chromosome could potentially be explained if DSB repair is delayed. With a fixed on/off rate for PRDM9, the longer DSBs persist, the more time MRE11 may have to process and remove SPO11-bound DNA, as PRDM9 will eventually exit the hotspot. Similarly, in PRDM9-deficient mice with defective repair, DSBs persist longer, and SPO11-RI is absent. However, an increased lifespan of resected DSBs is not always associated with the removal of SPO11-RI. For example, loss of ZCWPW1, a dual histone methylation reader of PRDM9, does not impact DSB placement or SPO11-RI formation ^48^. *Zcwpw1^−/−^* spermatocytes display decreased DSB repair and asynapsis phenocopying loss of PRDM9 ^48^; however, SPO11-RI is present at WT levels in *Zcwpw1^−/−^* spermatocytes. Thus, failure in synapsis and DSB repair does not necessarily lead to an increased fraction of breaks that are fully and symmetrically processed by MRE11.

We find limited evidence of SPO11-RI in ATM-null cells. Instead there is a concomitant increase in SPO11cc and MRE11 binding due to incompletely processed DSBs. Recent studies have demonstrated that some of these DSBs arising in *Atm*^−/−^ mice can be repaired by detrimental non homologous end-joining (NHEJ) (A. Lukaszewicz and M. Jasin, personal communication). Given the dramatic *in vitro* activity of TDP2 in *Atm*^−/−^ spermatocytes, SPO11cc might be rejoined in part by TDP2-dependent NHEJ *in vivo*. Direct hydrolysis of PRDM9-blocked SPO11 by TDP2 in WT cells would also likely result in aberrant end-joining. However, we found that purified TDP2 does not act on SPO11-RI *in vitro* (**Fig. 3d**). Although it remains unclear why SPO11-RI is an inefficient substrate for TDP2, it is possible that TDP2 cannot recognize SPO11-RI due to steric hindrance associated with the heteroduplex DNA. In any case, if TDP2 were similarly inactive on SPO11-RI *in vivo*, the formation of SPO11-RI could serve to prevent end-joining and instead favor repair of SPO11cc towards high fidelity HR.

In conclusion, both ATM and PRDM9 orchestrate mammalian SPO11-processing in a manner that influences meiotic DSB repair.

## Acknowledgements

We thank Todd Macfarlan, Mohamed Mahgoub, Michael Lichten, and Kevin Brick for comments on the manuscript; Francesca Cole, Todd Macfarlan, Mohamed Mahgoub, Scott Keeney, Agnieszka Lukaszewicz, Maria Jasin, Anjali Hinch and Petko Petkov for sharing unpublished data and insightful suggestions; Petko Petkov for PRDM9 KO mice and Scott Keeney for DMC1 knockout mice; Keith W. Caldecott for purified human TDP2; and Jennifer Wise and Kelly Smith for assistance with animal work. The A.N. laboratory is supported by the Intramural Research Program of the NIH, an Ellison Medical Foundation Senior Scholar in Aging Award (AG-SS-2633-11), the Department of Defense Idea Expansion (W81XWH-15-2-006) and Breakthrough (W81XWH-16-1-599) Awards, the Alex Lemonade Stand Foundation Award, and an NIH Intramural FLEX Award.

## Author contributions

J.P., W.W., and A.N. designed experiments; J.P., N.S., A.C., E.C, and A.D. performed experiments; R.A.D. provided purified proteins; S.Y. provided critical insight and interpretations; W.W., J.P., Y.M. and A.N. analyzed and interpreted the data; T.P. and A.N. supervised the research and provided advice. J.P., W.W. and A.N. wrote the manuscript with comments from the authors.

## Competing interests

The authors declare no competing interests.

---

**Extended Data Figure 1.**
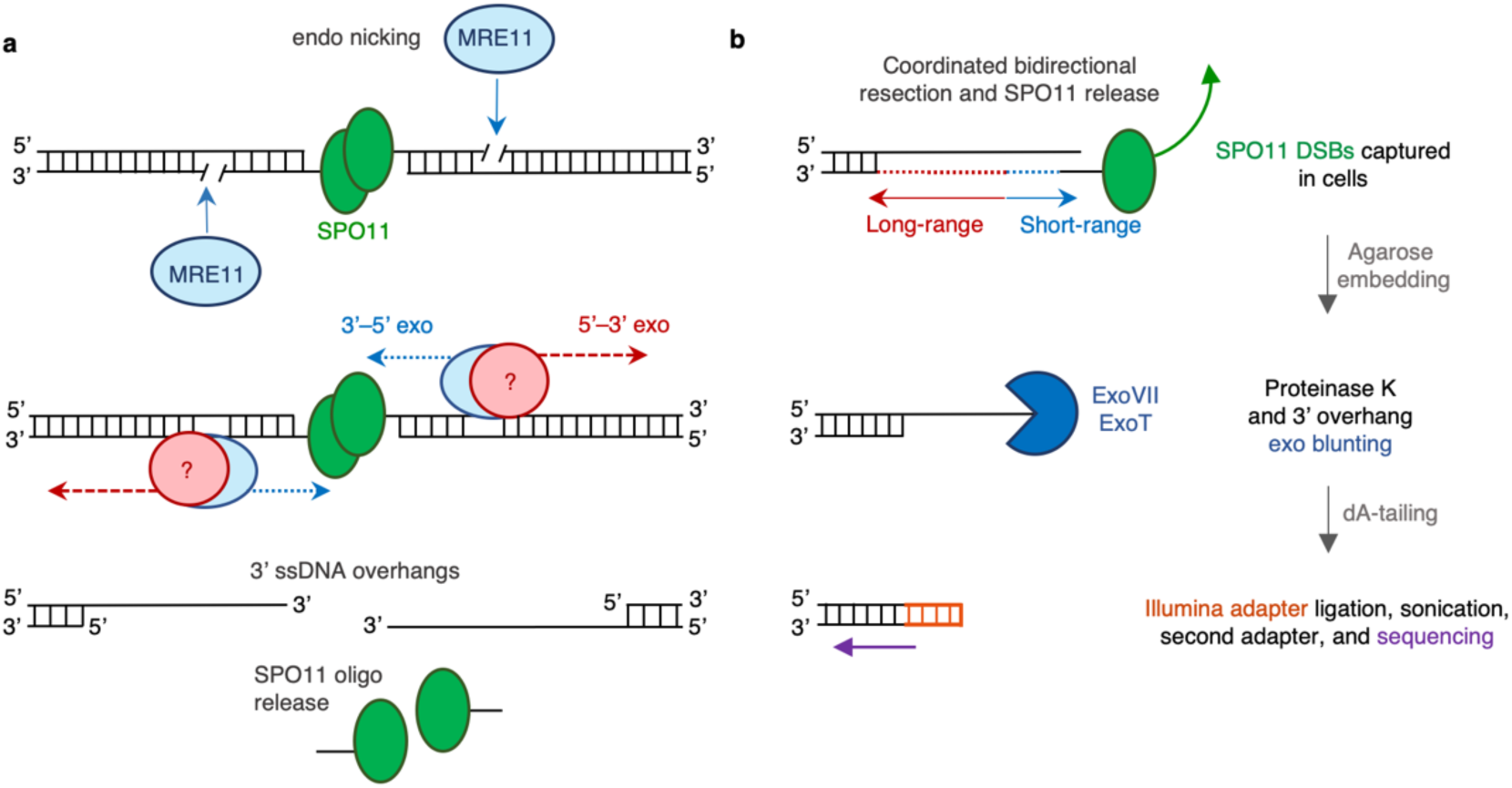
SPO11 generates meiotic DSBs that are resected and detectable by END-seq. (**a**) Illustration of meiotic break generation and processing. SPO11 induces a double-strand break (DSB) and remains covalently bound to both DNA ends. MRE11 recognizes the DSB and induces a nick on the SPO11-bound strand. Tightly coordinated short-range 3’-5’ resection by MRE11 and long range 5’-3’ resection by an unknown nuclease generates 3’ overhangs for homology search. MRE11 activity releases chromatin-bound SPO11 attached to short oligonucleotides (SPO11-oligos). (**b**) Schematic of END-seq detection of SPO11 DSBs (one side of the DSB is shown for simplicity). *In vivo* processing of SPO11 by coordinated bidirectional resection removes covalently bound SPO11 and produces a 3’ overhang present at the time of END-seq preparation and agarose embedding. Initial END-seq processing degrades all proteins by proteinase K and blunts ssDNA overhangs by nuclease digestion. Once fully blunted and dA-tailed, DNA ends are ligated to Illumina sequencing adapters, sheared, adapter ligated at other end of the fragment, and sequenced.

**Extended Data Figure 2.**
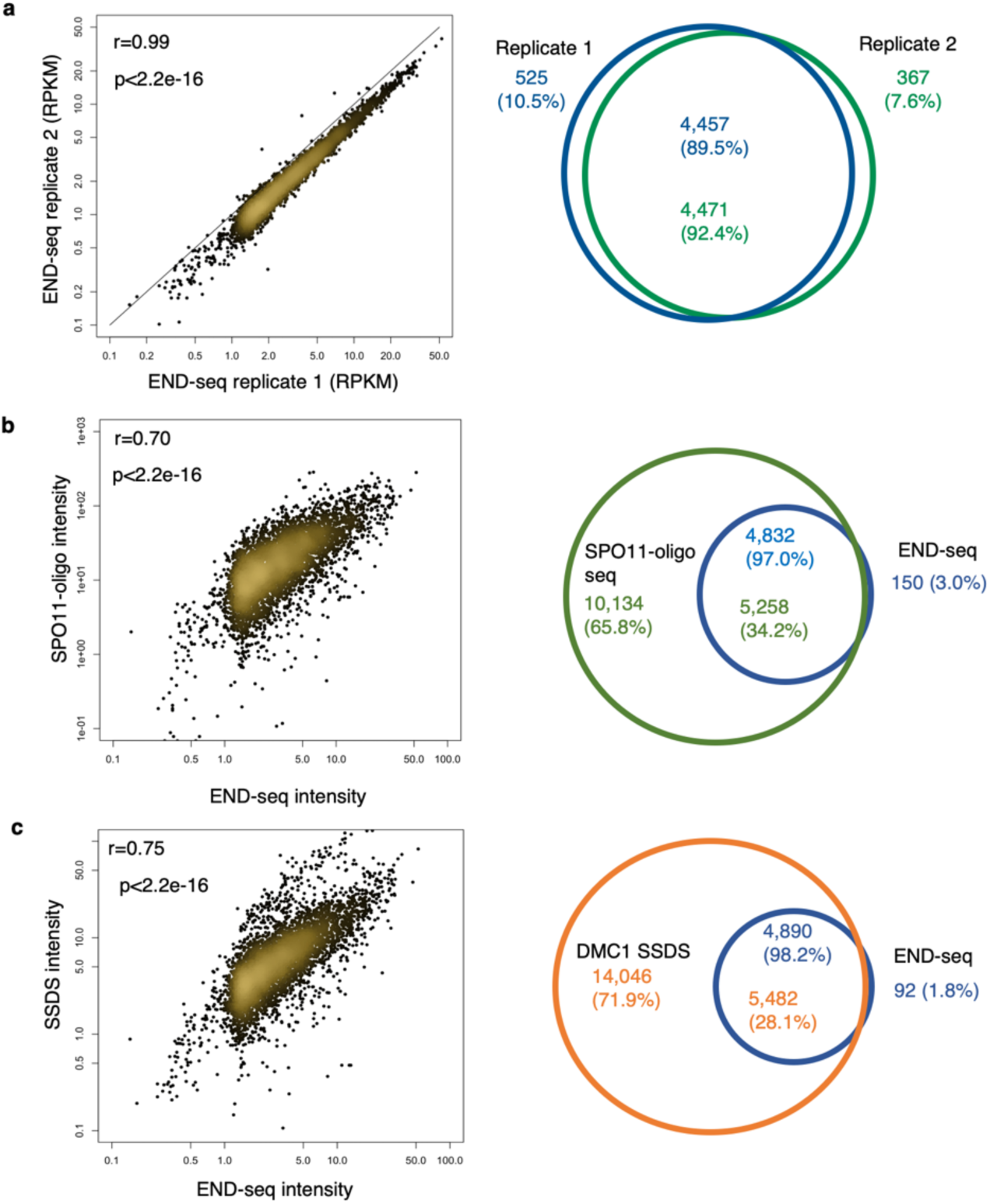
Comparisons of END-seq to previous hotspot mapping datasets. (**a**) Comparison of two WT END-seq biological replicates each made from 20 pooled juvenile mice. Left panel: Correlation (Pearson’s r) between break intensity of the two replicates in a ±3kb window around SPO11 summits. Right panel: Venn diagram showing overlap of peak calling from two END-seq replicates. P value <2.2e-16, fisher’s exact test. (**b**) Comparison of END-seq and SPO11-oligo sequencing hotspot mapping. Left panel: Correlation (Spearman’s r) of END-seq and SPO11-oligo intensity in a ±3kb window around SPO11 summits. Right panel: Venn diagram showing overlap of END-seq peaks and SPO11-oligo peaks. P value <2.2e-16, fisher’s exact test. (**c**) Comparison of END-seq and DMC1 SSDS hotspot mapping. Left panel: Correlation (Spearman’s r) of END-seq and SSDS intensity in a ±3kb window around SPO11 summits. Right panel: Venn diagram shows overlap of END-seq peaks and SSDS peaks. P value <2.2e-16, fisher’s exact test.

**Extended Data Figure 3.**
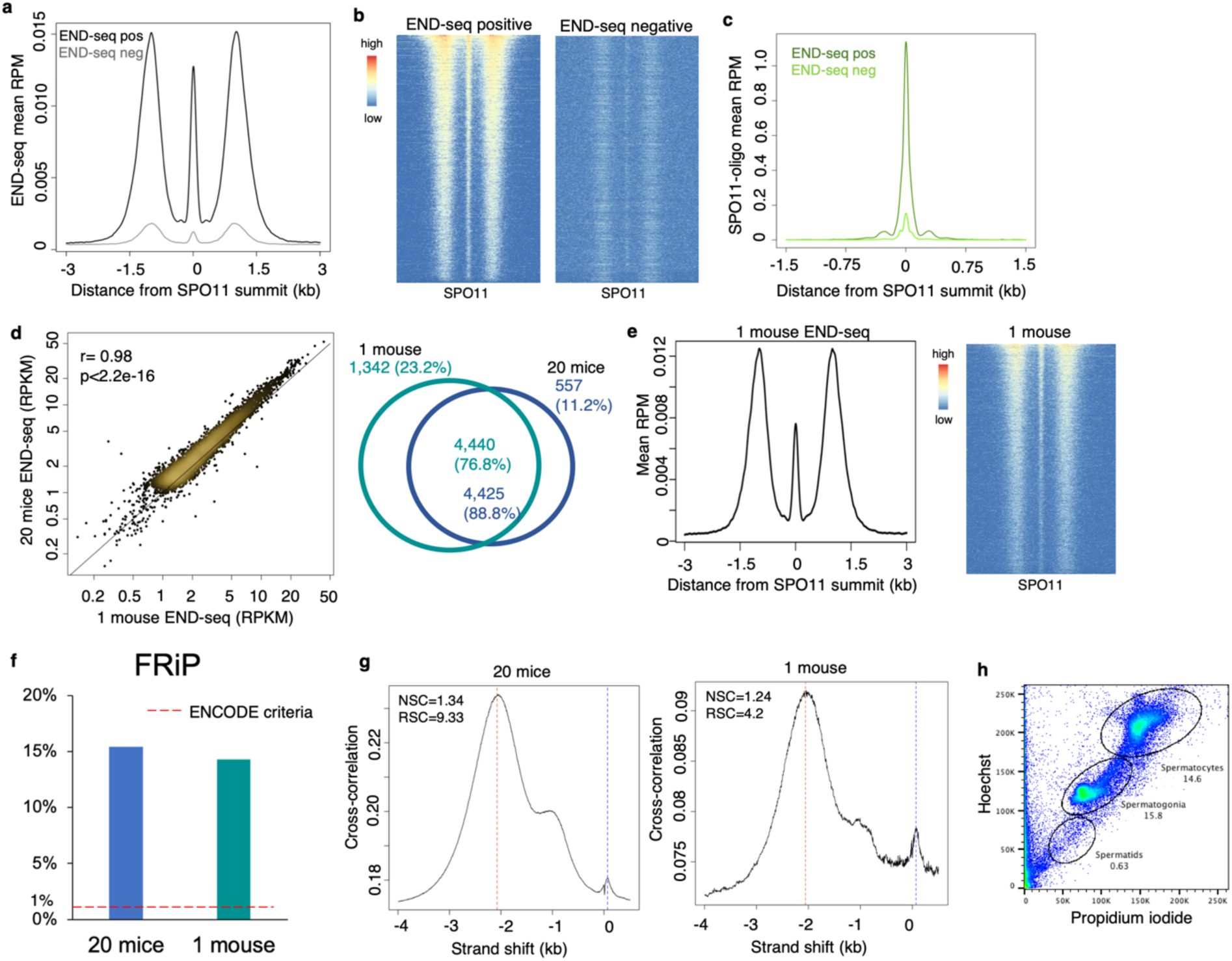
Analyses of END-seq sensitivity. (**a**) Aggregate plots of END-seq signal at SPO11-oligo sites for END-seq peak called hotspots (END-seq positive) versus non-peak called hotspots (END-seq negative) in a ±3kb window around the hotspot centers. (**b**) Heatmaps of END-seq signal for END-seq positive/negative hotspots in a ±2.5kb window around SPO11 summits. Break pattern is retained at all hotspots. (**c**) Aggregate plots of SPO11-oligo signal at END-seq positive versus END-seq negative hotspots in a ±1.5kb window around SPO11 summits. END-seq negative hotspots are also the coldest SPO11 hotspots. (**d**) END-seq detects meiotic DSBs with as low sample input as one mouse. Left panel: Correlation (Pearson’s r) between a library generated from twenty mice versus a library from one 12 dpp mouse in a ±3kb window around SPO11 summits. Right panel: Venn diagram showing high overlap of called peaks between twenty mice and one mouse. P value <2.2e-16, fisher’s exact test. (**e**) END-seq from a single mouse retains full break pattern at all 5000 strongest hotspots. Left panel: Aggregate plot of END-seq signal from one mouse in a ±3kb window around SPO11 summits. Right panel: Heatmap of END-seq signal from one mouse in a ±3kb window around SPO11 summits, ordered by total read count. (**f**) Similarity in FRiP values for twenty mice versus one mouse. Recommended ENCODE value denoted by dotted red line. (**g**) Cross-correlation plot profiles for twenty mice and one mouse END-seq. The plot shows Pearson cross-correlations (CCs, y-axis) of read intensities between the plus strand and the minus strand, after shifting minus strand (x-axis). There are two peaks, one corresponding to read length (CCread, blue dash line) and the other one corresponding to the fragment length (CCfrag,red dash line). Normalized strand coefficient (NSC) is CCfrag divided by minimal CC value (CCmin) and relative strand coefficient (RSC) is the ratio of CCfrag-CCmin divided by CCread-CCmin. NSC and RSC values are labeled. Higher NSC and RSC values mean more enrichment. ENCODE’s recommendation for ChIP-seq: NSC ≥1.05 and RSC ≥0.8. (**h**) Flow cytometric analysis of spermatocyte fraction in bulk testicular cells used for END-seq. Bulk cells were stained with Hoechst 33342 and propidium iodide to separate spermatogenesis progenitor populations. Spermatocytes, which harbor DSBs, represented ∼15% of the total bulk population.

**Extended Data Figure 4.**
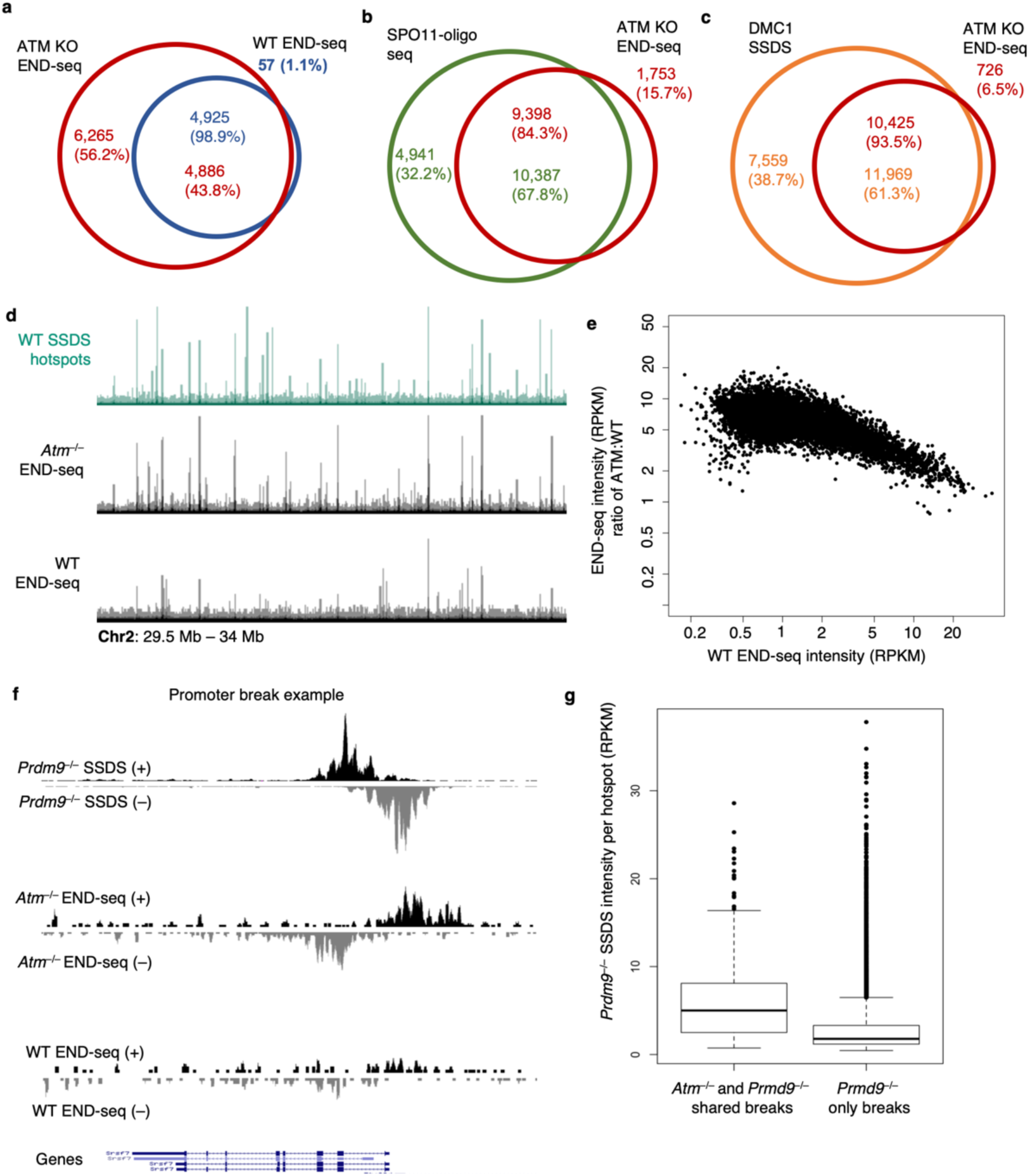
Comparisons of ATM-null END-seq to previous hotspot mapping methods. (**a**) Comparison of WT and *Atm*^−/−^ END-seq. Venn diagram shows overlap between WT and *Atm*^−/−^ END-seq peaks. P value <2.2e-16, fisher’s exact test. (**b**) Venn diagram shows overlap of *Atm*^−/−^ END-seq peaks and B6 SPO11-oligo sequencing peaks. P value <2.2e-16, fisher’s exact test. Of the 1,753 hotspots found in ATM-null END-seq and not SPO11-oligo seq, 48% (833 out of 1,753) overlap with breaks specific to *Prdm9*^−/−^ SSDS hotspots, of which 48% are at promoters. (**c**) Venn diagram shows overlap of *Atm*^−/−^ END-seq peaks and DMC1 SSDS peaks. P value <2.2e-16, fisher’s exact test. Of the 726 hotspots found in ATM-null END-seq and not SSDS, 82% (597 out of 726) overlap with breaks specific to *Prdm9*^−/−^ SSDS hotspots, of which 59% are at promoters. (**d**) Representative genome browser profiles of meiotic hotspots on Chromosome 2 for WT SSDS, *Atm*^−/−^ END-seq, and WT END-seq. Browser axis scales are equal for *Atm*^−/−^ and WT END-seq tracks to highlight the increased signal in the absence of ATM at typically cold hotspots. (**e**) The END-seq intensity of weaker hotspots preferentially increase more than stronger hotspots in *Atm*^−/−^ mice. WT RPKM per hotspot plotted against RPKM ratio of *Atm*^−/−^ to WT (signal normalized to spike-in control). (**f**) Genome browser example of an ATM-null END-seq break at a *Prdm9*^−/−^ SSDS promoter hotspot. Top (+) and bottom (–) strand-separated *Prdm9*^−/−^ SSDS and *Atm*^−/−^ END-seq tracks show significant signal at the promoter of the Srsf7 gene (bottom). WT END-seq tracks are shown to the same scale as ATM-null, illustrating that these breaks are specific to the loss of ATM regulation of SPO11. (**g**) Boxplot comparison of *Prdm9*^−/−^ SSDS intensity per hotspot of *Atm*^−/−^ and *Prdm9*^−/−^ shared hotspots versus hotspots found only in *Prdm9*^−/−^ mice. Hotspots shared between *Atm*^−/−^ END-seq and *Prdm9*^−/−^ SSDS are among the hottest *Prdm9*^−/−^ hotspots.

**Extended Data Figure 5.**
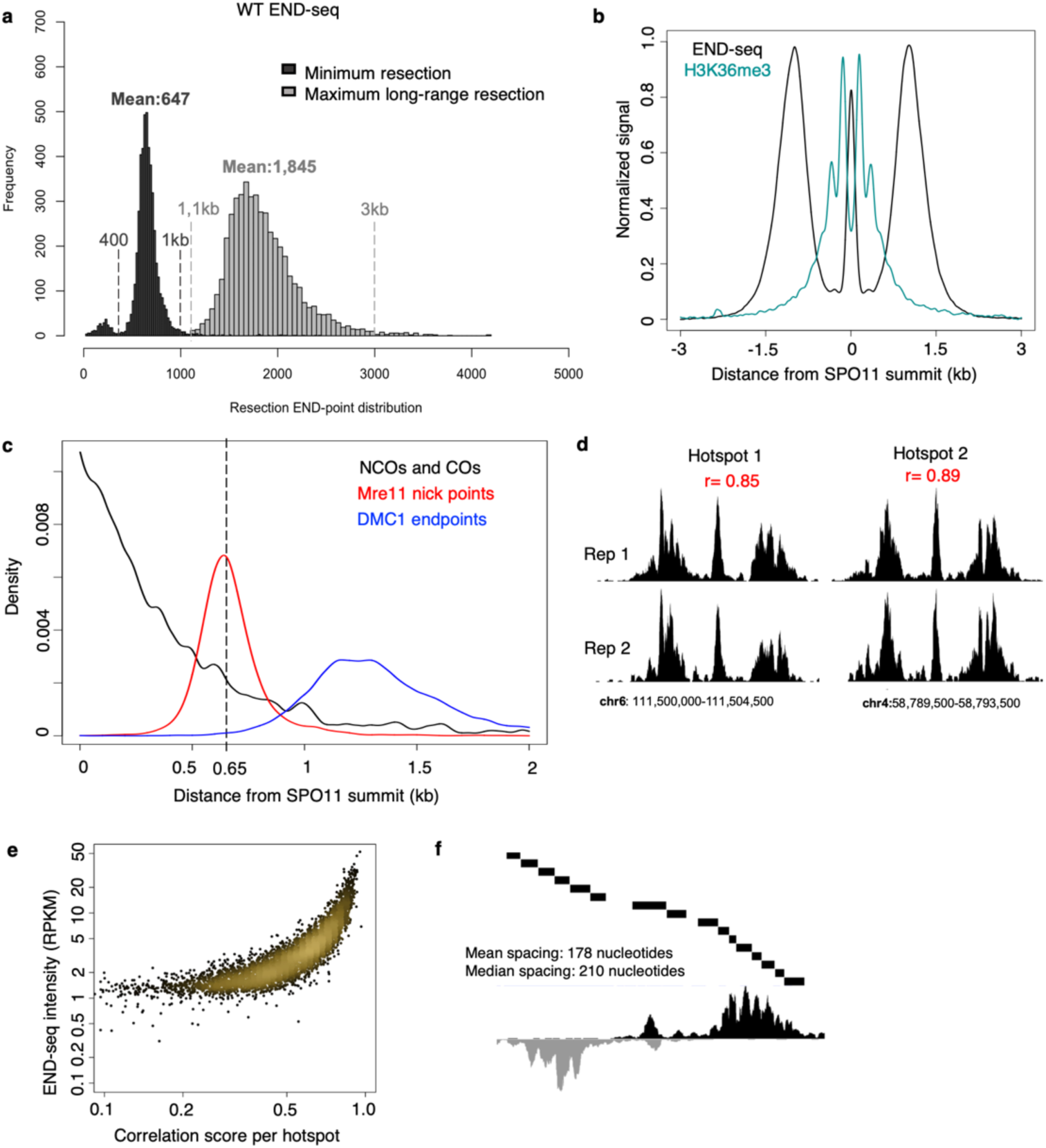
Resection length calculations and reproducibility likely reflect nucleosome occupancy. (**a**) Histogram distributions of END-seq minimum resection lengths and maximum long-range resection endpoints in WT spermatocytes. Mean values (bp) are listed. (**b**) Aggregate signal of WT END-seq and WT H3K36me3 ChIP-seq around SPO11-oligo summits at top 5000 breaks. Signals are normalized to the same height. (**c**) Distribution of short-range resection endpoints by END-seq (red), crossover breakpoints and noncrossover midpoints (black), and DMC1 resection endpoints (blue) relative to SPO11-oligo summits. (**d**) Between two biological END-seq replicates, hotspot break pattern is highly reproducible. Hotspots were binned into 20nt bins and reads per bin were correlated between replicates (Pearson correlation). Two hotspot examples demonstrate the reproducibility in resection pattern. (**e**) Correlation score (determined in “**d**”) plotted against END-seq intensity per hotspot. Hotter hotspots produce more reproducible resection patterns between replicates. (**f**) Example of subpeak calling within a single hotspot. Subpeak locations (top, 1 bar=1 subpeak) overlap well with END-seq resection pattern (bottom). Hotspot location is Chr1:68,484,499-68,495,00.

**Extended Data Figure 6.**
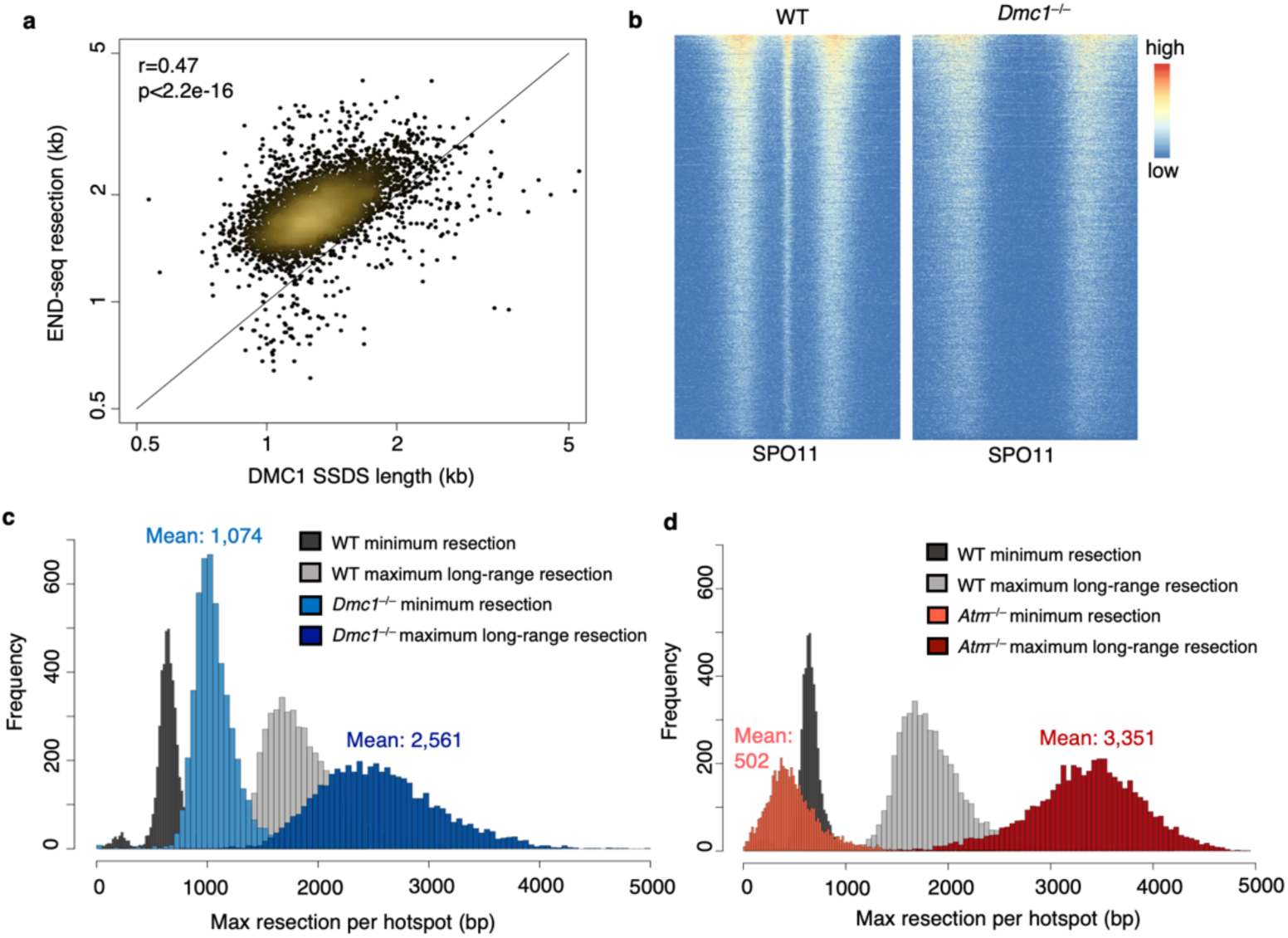
Increased resection lengths in DMC1- and ATM-null cells. (**a**) Correlation (Pearson’s r) of maximum long-range resection endpoints detected by END-seq and SSDS. (**b**) WT vs *Dmc1*^−/−^ END-seq heatmap of top 5000 breaks showing lack of central signal and increased short- and long-range resection in DMC1-null background. All hotspots show absence of central signal by heatmap in a ±2.kb window around SPO11 summits, ordered by total read count of WT END-seq. (**c**) Histogram distributions of WT and *Dmc1*^−/−^ END-seq short-range resection gap lengths and maximum long-range resection endpoints per top 5000 hotspots. Mean *Dmc1*^−/−^ values (bp) are listed. (**d**) Histogram distributions of WT and *Atm*^−/−^ END-seq short-range resection gap lengths and maximum long-range resection endpoints per top 5000 hotspots. Mean *Atm*^−/−^ values (bp) are listed.

**Extended Data Figure 7.**
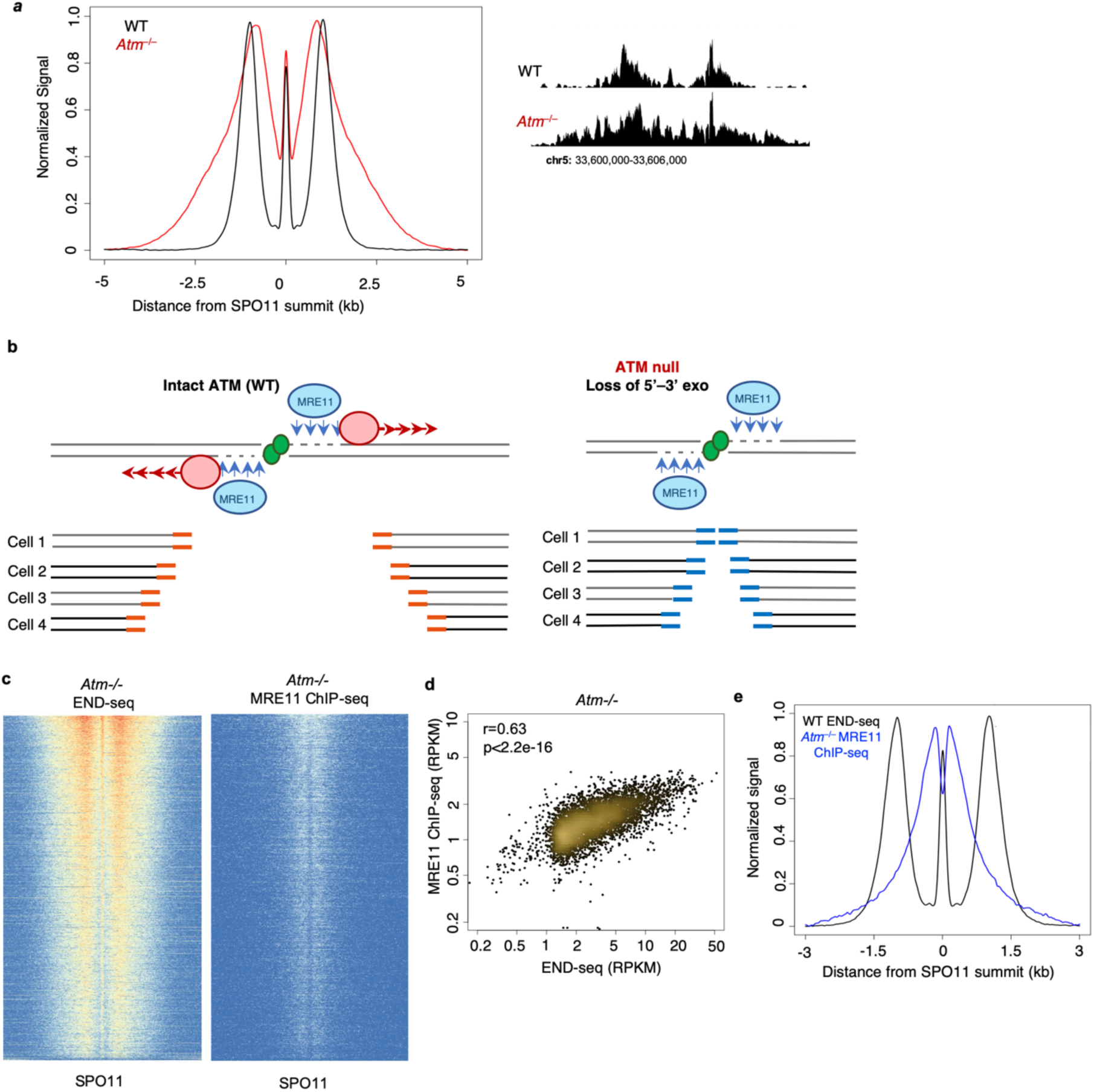
ATM coordinates resection machinery. (**a**) Left panel: Aggregate plots of END-seq signal in WT and *Atm*^−/−^ in top 5000 breaks around SPO11-oligo summits. Signals are normalized to the same height. Right panel: Single hotspot example of *Atm*^−/−^ resection pattern vs WT. (**b**) Illustration of ATM’s role in coordinating short- and long-range resection machineries. When ATM is intact (left), MRE11 3’-5’ resection and 5’-3’ long-range resection normally occur simultaneously, resulting in END-seq patterns with distinct boundaries between the read-less gap and long-range resection endpoints. In the absence of ATM (right), reads begin to accumulate within the typically read-less gap, likely indicating that a subset of DSBs undergo MRE11 endo nicking, yet do not engage long-range resection, resulting in adapter ligation (blue DNA segments) at the site of MRE11 nicking. (**c**) Heatmaps of END-seq and MRE11 ChIP-seq in *Atm^−/−^* around SPO11 summits. (**d**) Correlation (Spearman’s r) between END-seq intensity and MRE11 ChIP-seq per hotspot in *Atm^−/−^* spermatocytes. (**e**) Aggregate plot overlapping WT END-seq and ATM-null MRE11 ChIP-seq, normalized to the same height. MRE11 shows strong localization to the WT read-less gap.

**Extended Data Figure 8.**
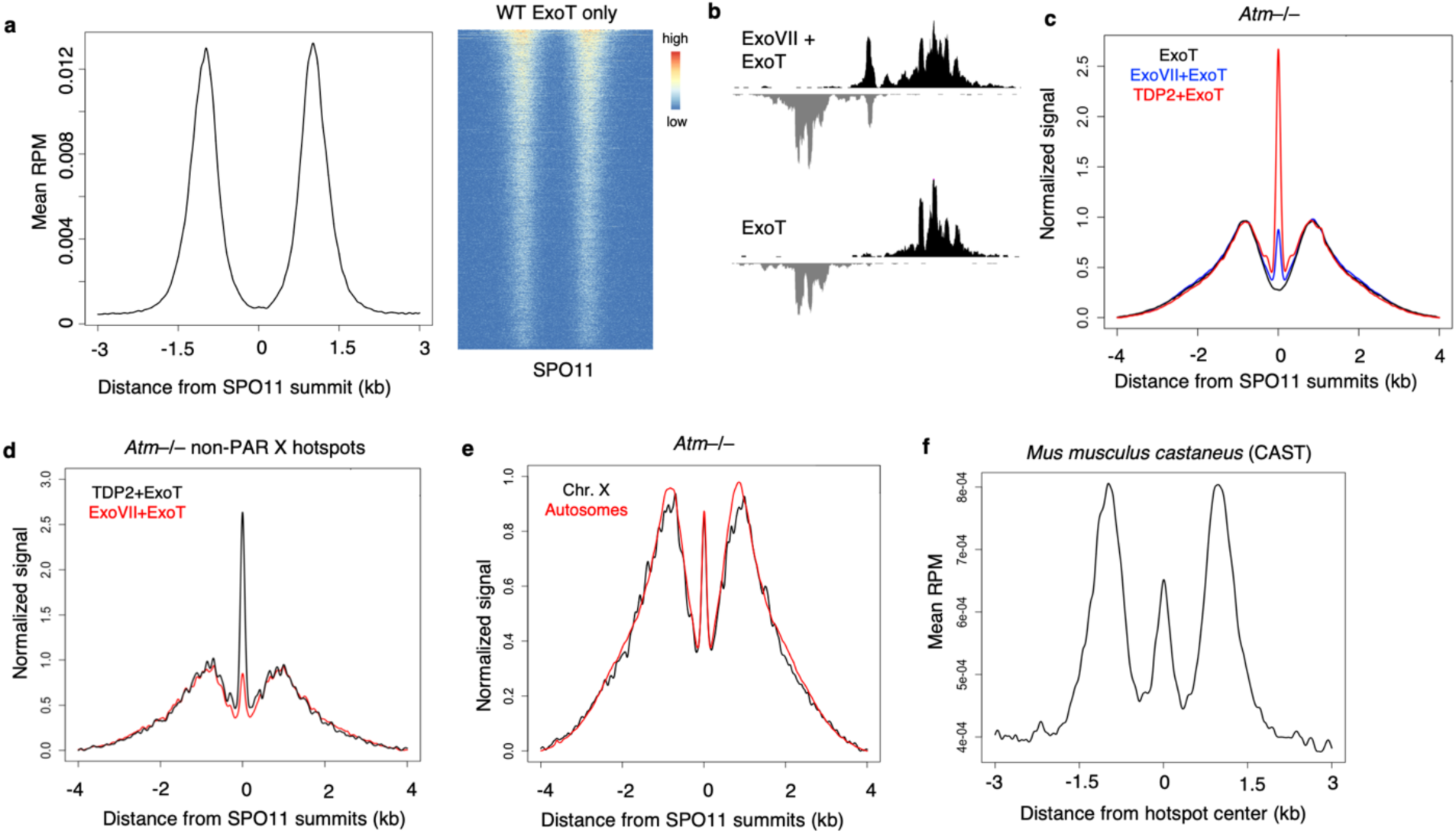
Detection of SPO11 cleavage complexes. (**a**) END-seq processing with ExoT blunting alone shows total absence of SPO11 central peak when signal is aggregated around SPO11-oligo summits (left). Heatmap of signal in ±3kb window around the hotspot centers, ordered by total read count of END-seq (right). (**b**) A single hotspot example of END-seq with ExoVII/ExoT versus with ExoT processing alone. SPO11 central peak detection entirely depends on ExoVII. (**c**) TDP+ExoT is more efficient at detecting SPO11cc than ExoVII+ExoT processing. Aggregate plot of *Atm^−/−^* END-seq signal (normalized to the same resection height) comparing ExoVII+ExoT, TDP2+ExoT, and ExoT alone. (**d**) SPO11cc is present at non-PAR X chromosome hotspots in the absence of ATM. Aggregate plots of *Atm^−/−^* END-seq signal (normalized to the same resection height) with TDP2+ExoT and ExoVII+ExoT. (**e**) Aggregate plot of *Atm^−/−^* END-seq signal (normalized to the same height) comparing SPO11cc intensity at non-PAR X chromosome hotspots and hotspots on all autosomes. (**f**) Aggregate plot of END-seq signal of an adult CAST parent from the B6xCAST F1 hybrid crosses in Fig. 4c. Unlike the hybrid pups in Fig. 4c, CAST males have prominent central signal at hotspot centers, determined by DMC1 SSDS.

**Extended Data Figure 9.**
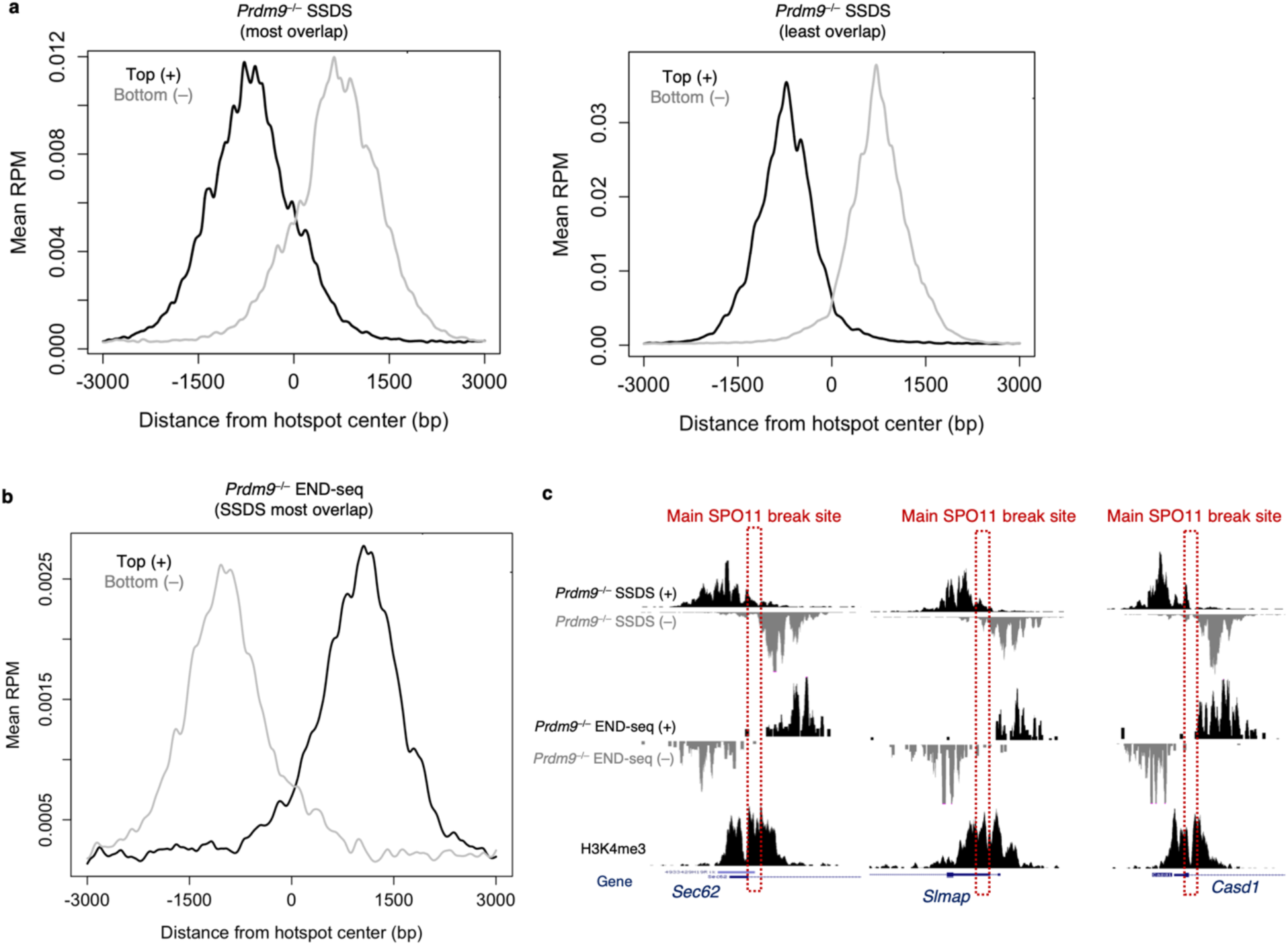
Characterization of PRDM9-independent hotspots. (**a**) Strand-separated aggregate plots of *Prdm9^−/−^* SSDS hotspots that have either the most top and bottom strand overlap (left, 375 hotspots) or least overlap (right, 199 hotspots), i.e. multiple SPO11 cut sites around the center (left) or a main SPO11 cut site (right). Overlap was determined by calculating, for each strand, the integration of signal from one side of the hotspot divided by the integration of total hotspot signal. “Most overlapped” hotspots have integration ratios of 0.75-0.85 (i.e. only 75-85% of total signal comes from one side of the hotspot) and “least overlapped” have ratios greater than 0.9. (**b**) Aggregated END-seq around *Prdm9^−/−^* SSDS hotspot center at sites described as “most overlapped” in “**a, left”.** At sites in which there is no clear center origin for SPO11 cutting, END-seq resection signal from multiple, adjacent cut sites within the promoter result in ambiguous central signal detection. We therefore focused analyses on the “least overlapped” hotspots described in “**a, right”**. (**c**) *Prdm9^−/−^* SSDS and END-seq tracks at additional default hotspots with minimal SSDS top and bottom strand overlap.

**Extended Data Figure 10.**
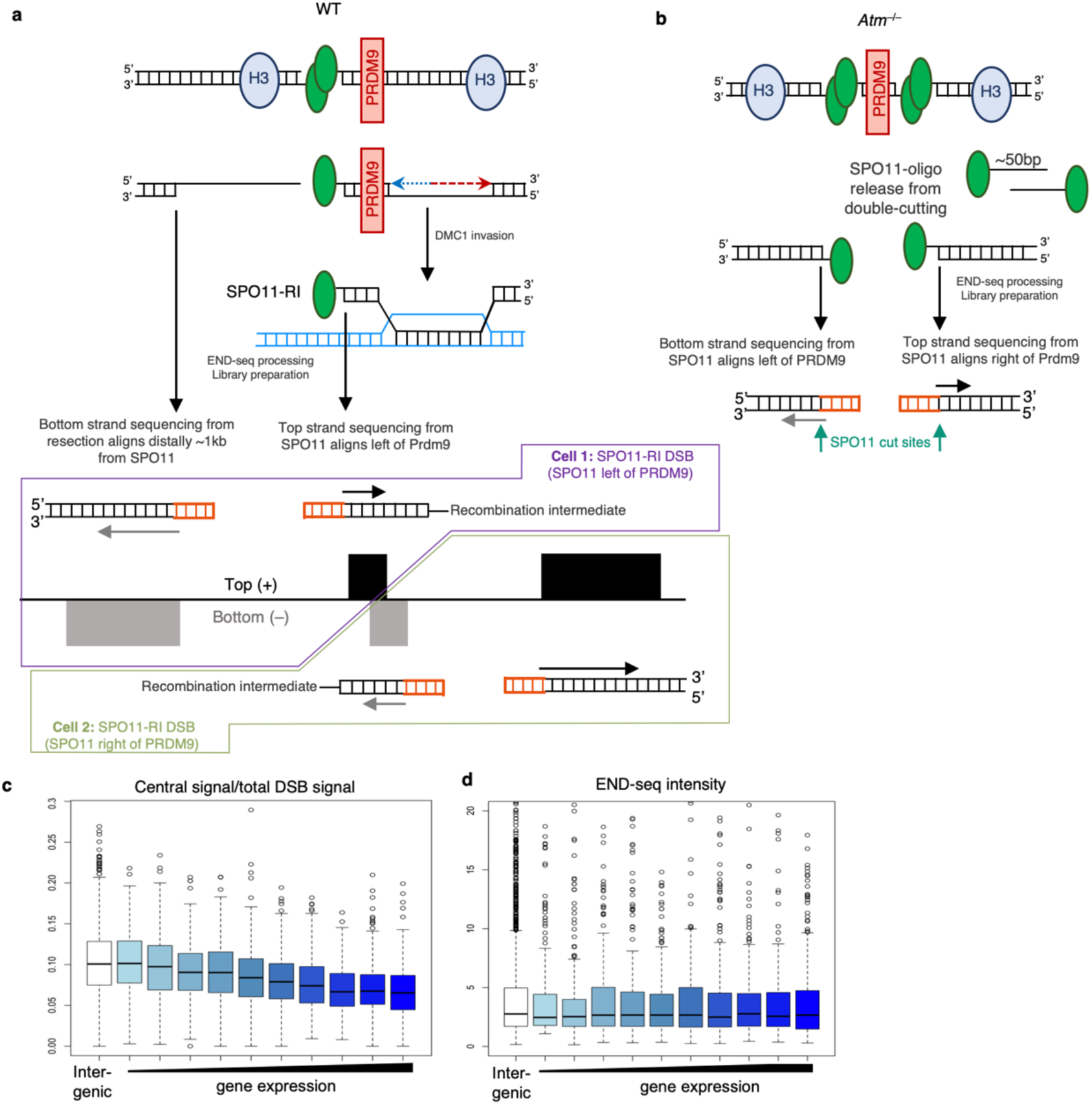
Models of WT and ATM-null SPO11 central signal origins and relationship with transcription. (**a**) Illustration of how bound PRDM9 acts as a barrier to SPO11 processing, resulting in a SPO11-RI structure at the center of hotspots (top). SPO11 cutting to one side of PRDM9 within the NDR may result in complete processing on one side of the break, generating a protein-free 3’ overhang, while MRE11 3’-5’ resection is blocked by PRDM9 on the other side of the break, leaving SPO11 covalently bound to a short stretch of dsDNA at the end of the DMC1-loaded ssDNA that extends as far as long-range resection traverses, ∼1kb from the NDR. Sequencing SPO11-RI generated from SPO11 that cut left of PRDM9 would result in top strand reads aligning to the left of the PRDM9 motif and bottom strand reads from fully resected ssDNA aligning at a great distance away from the break site (bottom illustration). Likewise, SPO11-RI generated from SPO11 that cut right of PRDM9 would result in bottom strand reads aligning to the right of the PRDM9 motif. An aggregation of both events within a population of spermatocytes gives strand separated pattern observed in “**a**” (bottom illustration). (**b**) Illustration of SPO11 double-cutting in *Atm^−/−^* cells that would show the correct polarity in top and bottom strands. Decreased MRE11 activity at these breaks would result in the direct sequencing of SPO11 cleavage complexes within the NDR. See also ^49^. (**c-d**) Boxplots comparing the ratio of WT END-seq central signal to total DSB END-seq signal per hotspot. Boxplots are categorized as either intergenic hotspots or genic hotspots (located within gene bodies) that are further categorized into ten groups based on gene expression data. Central signal decreases proportionally with increasing gene expression (**c**), while total break intensity per hotspots is unaffected by transcription (**d**).

## Methods

### Mice

C57BL/6 mice (Jackson Laboratory) were bred in-house, and male pups euthanized at 12-14 dpp for testes isolation. *Atm*^−/−^ mice (Barlow et al. 1996) were backcrossed eleven generations to C57BL/6 mice. *Exo1*^−/−^ mice were a gift from Winfried Edelmann. *Dmc1*^−/−^ mice were a gift from Scott Keeney. *53bp1*^−/−^ mice were a gift from Junjie Chen. *Prdm9*^−/−^ mice were a gift from Petko Petkov. *Brca1^+/△^*^11^ mice were obtained from the NCI mouse repository. *53bp1^S25A^* mice were generated as described ^34^. *p53*^−/−^ mice were purchased from Taconic Biosciences. Strains were crossed to generate *Brca1^Δ11^53bp1*^−/−^, *Brca1^△11^53bp1^S25A^*, and *Brca1^△11^p53^+/–^* mice. CAST/EiJ (males) were purchased from Jackson Laboratory and were crossed with C57BL/6 (females) for one generation to produce B6xCAST F1 hybrids, respectively. All mouse breeding and experimentation followed protocols approved by the National Institutes of Health Institutional Animal Care and Use Committee.

## Method details

### Mouse testicular cell isolation

Adapted from (Baker et al. 2014). Testes were dissected from 12-14 dpp male mice and placed into a 6 cm tissue culture dish containing DMEM. Tunica albuginea were removed under a microscope, and tubules were gently dissociated with forceps and placed into 50 mL tube containing 20 mL DMEM. After tubules settled to the bottom of the tube, DMEM was aspirated and replaced with 20 mL DMEM containing 0.5 mg/mL Liberase TM (Roche, 5401127001) and incubated at 32°C for 15 min at 500 rpm. Tubules were washed once with fresh DMEM, replaced with 20 mL DMEM containing 0.5 mg/mL Liberase TM and 100 U DNase I (ThermoFisher, EN0521), and incubated at 32°C for 15 min at 500 rpm.

Tubules were disrupted by gentle pipetting and passed through a 70 μm Nylon cell strainer (Falcon) repeatedly until tissue debris was fully removed. Cells were pelleted at 1500 rpm at 4°C for 5 min and washed with 10 mL DMEM. Cells were filtered through a 40 μm Nylon cell strainer (Falcon) repeatedly until debris was fully removed and pelleted at 1500 rpm at 4°C for 5 min. Cells were resuspended in 1 mL PBS and counted.

### END-seq

#### Embedding cells into agarose plugs

Single-cell suspensions of bulk testicular cells were immediately embedded after isolation into 0.75% agarose plugs. After isolation, cells in 1 mL PBS were diluted to 5-7 million bulk cells/mL PBS and separated into 1 mL of cells per 1.5 mL tube for plug making. Spike-in cells were added at 2% of bulk cell number per tube/plug. Multiple plugs were made per sample if necessary, depending on number of mice and total cell number isolated, processed in the same tube, and DNA later combined after plug melting.

A detailed description for embedding cells into agarose plugs and general END-seq procedure can be found in (Canela et al. 2017, Canela et al. 2019). Briefly, agarose embedded cells were immediately lysed and digested with Proteinase K after agarose solidification. Plugs were then washed with TE, treated with RNase, and stored at 4°C for no longer than one week before the next series of enzymatic reactions.

#### Enzymatic reactions

Unless otherwise noted, plugs were treated with sequential combination of Exonuclease VII (NEB) for 1 hr at 37°C followed by Exonuclease T (NEB) for 45 min at 24°C to blunt DNA ends before Illumina adapter ligation (Canela et al. 2019). For experiments in which only Exonuclease T was used to blunt, plugs were digested with Exonuclease T for 1 hr at 24°C. For experiments with purified human TDP2 (gift from Keith W. Caldecott), plugs were treated with 500 pM TDP2 (in 50 mM Tris-HCl pH 8.0, 10 mM MgCl2, 80 mM KCl, 1 mM DTT, and 0.05% Tween-20) for 4 hr at 24°C, followed by Exonuclease T digestion for 1 hr at 24°C.

For experiments with purified human MRE11/RAD50/NBS1 (MRN) and CtIP (Tanya Paull lab), plugs were treated with 50 nM MRN (MR and NBS1 were pre-incubated for 10 min at 4°C prior to reactions) with or without 80 nM CtIP (in 25 mM MOPS pH 7.0, 20 mM Tris-HCl pH 8.0, 80 mM NaCl, 1 mM DTT, 1 mM ATP, 5 mM MgCl2, 1 mM MnCl2, and 0.2 mg/mL BSA) for 1.5 hr at 37°C. Plugs were treated again with proteinase K for 1 hr at 50°C to degrade any remaining bound protein, followed by Exonuclease VII and Exonuclease T digestion as previously described.

For all enzymatic reactions, subsequent steps of A-tailing, adapter ligation, plug melting, chromatin shearing, and second round of adapter ligation for sequencing were performed exactly as previously described (Canela et al. 2017, Canela et al. 2019).

### ChIP-seq

10 million bulk testicular cells were fixed in 1% formaldehyde (Sigma, F1635) at 37°C for 10 min. Fixation was quenched with glycine (Sigma) at a final concentration of 125 mM. Cells were washed twice with cold PBS, and pellets were snap frozen on dry ice and stored at −80°C until sonication. Sonication, immunoprecipation, and library preparation were performed as previously described in (Canela et al. 2019). Briefly, frozen pellets were resuspended in 1 mL cold RIPA buffer (10 mM Tris-HCl pH 7.5, 1 mM EDTA, 0.1% SDS, 0.1% sodium deoxycholate, 1% Triton X-100, and 1 Complete Mini EDTA-free proteinase inhibitor tablet (Roche)) and sonicated using a Covaris S220 at duty cycle 20%, peak incident power 175, and cycle/burst 200 for 30 min at 4°C.

Chromatin was precleared with 40 μL prewashed Dynabeads Protein A (ThermoFisher) for 30 min at 4°C, followed by incubation with 40 μL Dynabeads Protein A bound to either 6 μL anti-H3K4me3 (Millipore, 07-473) or 4 μL anti-MRE11 (Novus, NB100-142) overnight at 4°C. Beads were washed and crosslinking reversed the next day as described in (Canela et al. 2019). Immunoprecipitated DNA was removed from beads and stored at −20°C until library preparation, which was performed as described in (Canela et al. 2019).

### END-seq Data Analysis

#### Mapping

Reads were aligned to the mouse (GRCm38p2/mm10) genome using Bowtie version 1.1.2 ^50^ and 3 mismatches were allowed and the best strata for reads were kept with multiple alignments (-n 3 -k 1 -l 50). Functions ‘‘view’’ and ‘‘sort’’ of samtools (version 1.6) ^51, 52^ were used to convert and sort the mapping output to sorted bam file.

#### Peak calling

Peaks were called using MACS 1.4.3 ^53^. As the strand separated aggregate plots (Figure 2a) show ∼2kb distance, END-seq peaks were called using the parameters: --shiftsize=1000, -- nolambda, --nomodel and --keep-dup = all. Peaks with >2.5 fold-enrichment are kept and those within blacklisted regions (https://sites.google.com/site/anshulkundaje/projects/blacklists) were filtered.

#### Total Break Number Estimation

For estimation of total breaks and comparison between genotypes, we added a spike-in control into END-seq samples which consists of a G1-arrested Ableson-transformed pre-B cell line (*Lig4^-/-^*) carrying a single zinc-finger-induced DSB at the TCRβ enhancer. This site is expected to break in all spike-in cells, which were mixed in at a 2% frequency with bulk testicular cells. END-seq signal was calculated, as RPKM, within ±5kb window around all hotspot centers. Total intensity was divided by the signal around the spiked-in breaks and then divided by 50 since the spiked-in was added at a 1:50 ratio (2%).

#### Subpeaks Identification

As the pattern of END-seq peaks are consisted of continuously sharp peaks and it is quite reproducible between experiments and to estimate the distance between the shar peaks and study its relation with nucleosome positioning, END-seq peaks were split as subpeaks by PeakSplitter tool of PeakAnalyzer with default parameters (https://www.bioinformatics.org/peakanalyzer/wiki/Main/Overview, ^54^) and the distances of sharp peak summits within each break were calculated.

#### Signal to Noise Analysis

ENCODE measures Signal-to-noise(S/N) ratio by fraction of reads in peaks (FRiP) and cross-correlation profiles (CCPs) for ChIP-seq. We used it for END-seq here.

The FRiP value is fractions of reads that mapped into called peaks without any filtering. The higher FRiP, the more enrichment. ENCODE recommends the threshold for FRiP is more than 1% for ChIP-seq.

The CCPs assess the quality of END-seq enrichment over background independent of peak calling. The plot shows the Pearson cross-correlations (CCs, y-axis) of reads intensities between the plus strand and the minus strand, after shifting minus strand (x-axis). There are two peaks, one corresponding to read length (CCread, blue dash line) and the other one corresponding to the fragment length (CCfrag, red dash line). Normalized strand coefficient (NSC) is CCfrag divided by minimal CC value (CCmin) and relative strand coefficient (RSC) is the ratio of CCfrag-CCmin divided by CCread-CCmin. The higher NSC and RSC, the more enrichment. ENCODE recommends an NSC ≥1.5 and RSC ≥0.8 for ChIP-seq.

The CCPs profile, NSC value and RSC value were generated by Phantompeakqualtools (https://github.com/kundajelab/phantompeakqualtools, ^21, 55^). Different to ChIP-seq pattern which plus strand signal is at the left of minus strand signal, END-seq has opposite direction, so the shift size of the fragment is negative. Unsurprising, the absolute shift values are consistent with the distance of two long-range resection peaks showed by the aggregate plots.

#### Quantification of short-range resection, mean long-range resection and maximum long-range resection

Short-range resection was measured by a sliding window containing ten 10bp bins, starting from the peak of long-range resection signal into the hotspot center. Mean value of ±500bp window around the peak site was used to estimate the normal distribution of the resection endpoints. Comparing the real signal of the sliding window to it, when the window got p-value ≤0.05 by one-sample t-test (alternative=”less”), the distance from hotspot center to the window midpoint was defined as the short-range resection length. Plus strand and minus strand were calculated separately.

For each hotspot, the long-range resection describes the distribution of resection endpoints among the cell population. Mean long-range resection was quantified based on the definition of expect value of a distribution as:

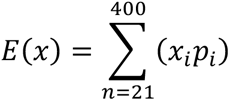

RPM were calculated for 10-bp bins in a ±4kb window of each SPO11 summit. The first bins were not considered, the calculation started from the 21^th^ bin. *x_i_* is distance of the current bin to SPO11 summit and *p_i_* is the probabilities calculated as RPM of the current bin / total RPM from 21^th^ to 400^th^ bins. Plus strand and minus strand were calculated separately. Top 250 breaks were used for the boxplot of all mean resection comparison.

To measure maximum long-range resection, a sliding window containing twenty 20-bp bins was used, starting from the peak of long-range resection to further region until more than half of bins within current window had lower intensity than background. The location of the last bin with a detectable signal over background is regarded as the maximum resection endpoint. Dynamic background was determined by the maximum intensity of 20-bp bins within 5kb∼6kb region away from SPO11 summit for individual hotspot. Plus strand and minus strand were calculated separately.

#### Estimation of SPO11-RI and unresected SPO11cc

The fraction of central signal detected by ExoVII+ExoT or TDP2+ExoT in B6 was calculated using the total area substrate the area of ExoT blunting only after forcing them to the same height and then divided by ExoVII+ExoT total area. We got 11% for ExoVII+ExoT and 1.7% for TDP2+ExoT. As the central signal of ExoVII+ExoT consists of both SPO11-RI and unresected SPO11cc, we used TDP2 to measure fraction of the unresected SPO11cc. The detection ability of ExoT for the small overhangs of SPO11cc and long overhangs of SPO11-RI may differ. Assuming ExoT detected unresected SPO11cc and SPO11-RI at the same level, the unresected SPO11cc fraction will be 1.7%. However, ExoT might work better for unresected SPO11cc detection than SPO11-RI and assume it captures all unresected SPO11cc. We firstly quantified the total break number for ExoT blunting sample using spike-in normalization which is 93. Then multipled the ratio of the central signal with TDP2+ExoT vs ExoT only (∼2%). The number of unresected SPO11cc is ∼2 and is 0.6% of total break number. Thus, the fraction of unresected SPO11cc could be 0.6% ∼ 1.7% and SPO11-RI is 9.3% ∼ 10.4%.

#### Mouse Hybrid END-seq analysis

Single nucleotide polymorphisms (SNP) were annotated for CAST genome by the Sanger Mouse Genome Project ^56^ based on B6 mouse reference genome (mm10). Only SNPs (n=20,668,274) passed all filter criteria were employed to generate an “N-masked” genome. 14,951 hotspots in B6xCAST hybrid genome were download from GSE75419.. Since Bowtie doesn’t allow gapped alignment for Ns, the hybrid END-seq reads were aligned by Bowtie2 version 2.3.5.1 ^57^ with options -N 1 -k 1.

### SPO11-oligos, SSDS and ChIP-seq and RNA-seq Data Analysis

#### SPO11-oligos

Reads were aligned to the mouse (GRCm38p2/mm10) genome using Bowtie version 1.1.2 with options -n 2 -m 1 -l 50. Peaks were called using MACS 1.4.3. Peaks were called using the parameters: --shiftsize=73, --nomodel and then filtered by fold-enrichment (FC ≥10) as well as the blacklists. SPO11 summits from SPO11-oligo seq overlapped the B6 END-seq peaks (∼5000) were used for most of the aggregate plots except the B6xCAST hybrid data.

#### DMC1 SSDS

Type1 alignments and hotspots’ coordinates of SSDS data were directly download from GEO database. B6xCAST hybrid SSDS was analyzed as described in END-seq part.

#### H3K4me3 and H3K36me3 ChIP-seq

For our H3K4me3 ChIP-seq and the public H3K36me3 ChIP-seq, reads were aligned to the mouse (GRCm38p2/mm10) genome using Bowtie version 1.1.2 with options -n 2 -m 1 -l 50.

#### RNA-seq

For RNA-seq, reads were aligned to the mouse (GRCm38p2/mm10) genome using TopHat version 2.1.1 (PMID: 19289445) with default parameters. The gene expression level of RPKM were determined by Cuffnorm version 2.2.1 ^58^ based on the annotation from RefSeq. Mean values across samples were used.

### Data Visualization

Aligned-reads bed files were first converted to bedgraph files using bedtools genomecov (PMID: 20110278) following by bedGraphToBigWig to make a bigwig file ^59^. Visualization of genomic profiles was done by the UCSC browser ^60^. Genome browser profiles were normalized to the library size (RPM). Heatmaps were produced using the R package ‘pheatmap’.

Aggregate plots for sequencing data around hotspot centers or SPO11 summits were performed as follows: A fixed window was defined around all sites genome wide. The number of reads overlapping each nucleotide was calculated. The aggregate signal was smoothed using smooth.spline function in R. For END-seq, only the adapter ligated endpoints were used.

### Statistical Analysis

Statistical analysis was performed using R version 3.5.0 (http://www.r-project.org). The statistical tests are reported in the figure legend and main text.

### Public data used

Public data used:
| **Data type** | **GEO accession** | **Reference** |
| --- | --- | --- |
| RNA-seq | GSE61613 | 61 |
| H3K36me3 ChIP-seq | GSE76416 | 62 |
| Prdm9<sup>-/-</sup> SSDS | GSE99921 | 24 |
| Hybrid, B6 and CAST SSDS | GSE75419 | 20 |
| SPO11-oligo seq | GSE84689 | 8 |
| Crossovers and noncrossovers | SRP189007 | 15 |

